# A complex containing lysine-acetylated actin inhibits the formin INF2

**DOI:** 10.1101/509133

**Authors:** A Mu, Tak Shun Fung, Arminja N. Kettenbach, Rajarshi Chakrabarti, Henry N. Higgs

## Abstract

INF2 is a member of the formin family of actin assembly factors. Dominant mis-sense mutations in INF2 link to two diseases: focal segmental glomerulosclerosis (FSGS), a kidney disease; and Charcot-Marie-Tooth disease (CMTD), a neuropathy. All disease mutations map to the autoinhibitory Diaphanous Inhibitory Domain (DID). Curiously, purified INF2 is not autoinhibited, suggesting the existence of additional cellular inhibitors. We purified an INF2 inhibitor from mouse brain, and identified it as a complex between lysine-acetylated actin (KAc-actin) and cyclase-associated protein (CAP). Inhibition of INF2 by CAP/KAc-actin requires INF2 DID. Treatment of CAP/KAc-actin with histone deacetylase 6 (HDAC6) releases INF2 inhibition, while HDAC6 inhibitors block cellular INF2 activation. INF2 disease mutants are poorly inhibited by CAP/KAc-actin, suggesting that FSGS and CMTD result from reduced CAP/KAc-actin binding. This is the first demonstrated role for lysine-acetylated actin: regulation of an actin assembly factor by a novel mechanism, which we call facilitated auto-inhibition.

## INTRODUCTION

Most cellular processes are controlled by regulation of key proteins, with two common regulatory mechanisms being autoinhibition and post-translational modification (PTM). In autoinhibition, a protein’s effector domains are sequestered by a cis-acting element. Many well-known regulatory proteins are autoinhibited^1, 2^, and require activation mechanisms to release the autoinhibitory interaction. PTM by small chemical groups (phosphorylation, acetylation, methylation), lipids (palmitoylation) or proteins (ubiquitinylation, SUMOylation) can trigger major changes in protein activity or location. Frequently, both autoinhibition and PTM serve to regulate the same protein^1, 3^.

Members of the formin family of actin assembly factors are known or predicted to be autoinhibited^4-6^. The 15 mammalian formins are key players in diverse cellular functions, including cytokinesis, cell motility, phagocytosis and organelle dynamics. A sub-set of formins (10 mammalian formins^7-9^) contains two auto-regulatory sequences, the N-terminal Diaphanous Inhibitory Domain (DID) and the C-terminal Diaphanous Auto-regulatory Domain (DAD). The auto-inhibitory mechanism of one formin, mDia1, has been described in detail, with high-affinity DID/DAD interaction inhibiting the ability of the Formin Homology 2 (FH2) domain to stimulate actin polymerization^10^. Binding of GTP-charged RhoA to mDia1’s GTPase Binding Domain (GBD) activates the formin by disrupting the DID/DAD interaction^11-13^. Atomic structures have been solved of both DID alone and DID/DAD complex, with a single DID point mutation in the DAD binding site being shown to eliminate autoinhibition^12^, ^14^.

Inverted Formin 2 (INF2) is a mammalian formin containing DID and DAD sequences. Biochemically, INF2 has multiple effects on actin^15-17^, as well as having both direct and indirect effects on microtubules^18-21^. In cells, INF2-mediated actin polymerization plays roles in mitochondrial fission, ER-to-mitochondrial calcium transfer, an endocytic vesicle trafficking, and Golgi structure^22-27^.

Dominant mis-sense mutations in INF2 are linked to two diseases, focal segmental glomerulosclerosis (FSGS), a kidney disease^28^; and Charcot Marie Tooth disease (CMTD), a peripheral neuropathy^29^. Interestingly, all of the >35 disease-linked mutations map to the DID, suggesting an important regulatory function for this domain. However, few disease mutations are in the DAD binding site. In addition, three results suggest that INF2 does not engage in canonical autoinhibition. First, INF2’s DID/DAD interaction is >10-fold weaker than that of mDia1^15^, ^30^, ^31^. Second, actin monomers bind to INF2’s DAD, and compete with DID^30^. Third, purified INF2 is constitutively active in biochemical actin polymerization assays^30^, in stark contrast to the strong autoinhibition of purified mDia1^32^. While these results suggest that INF2 is not regulated by canonical autoinhibition, cellular studies show that INF2 is tightly regulated in a manner dependent on DID/DAD interaction. Over-expression of wild-type INF2 causes no apparent change in cellular actin polymerization, while over-expression of a DID mutant that disrupts DID/DAD interaction causes unregulated actin polymerization^25^, ^30^.

These results suggest that an additional cellular factor is required to strengthen INF2’s DID/DAD interaction. In this work, we identify this inhibitory factor as a complex between cyclase-associated protein (CAP) and lysine-acetylated actin. Deacetylation of the CAP/actin complex by histone deacetylase 6 (HDAC6) abrogates inhibition. Small molecule inhibitors of HDAC6 block INF2 activation in cells. INF2 mutants linked to FSGS and CMTD are not inhibited by CAP/actin, suggesting a mechanism for disease.

## RESULTS

### INF2 is constitutively active in vitro but inhibited by DID/DAD interaction in cells

INF2 has two splice variants that differ in C-terminal sequence: INF2-CAAX is prenylated and tightly bound to ER^23-25^, while INF2-nonCAAX is not prenylated and localizes to cytosol^22, 26, 27^. To test regulatory mechanisms, we used INF2-nonCAAX due to its solubility in aqueous solution. We expressed human full-length INF2-nonCAAX (INF2-FL, Fig. 1a) in mammalian HEK293 cells as an N-terminally tagged fusion with GFP and Strep-tag II (GFP-Strep), and purified the protein to apparent homogeneity either with or without the GFP-Strep moieties (Fig. 1b). By sedimentation velocity analytical ultracentrifugation, INF2 is a dimer (Fig. 1c). Next, we examined the effect of purified INF2 on actin dynamics, comparing the full-length protein to a truncated INF2 construct which lacks the DID (INF2-FFC). Similar to INF2 purified from insect cells^30^, HEK293-purified INF2 accelerates actin polymerization from monomers with equal potency to INF2-FFC (Fig. 1d). These results show that INF2 is not autoinhibited in the purified state.

**Figure 1.**
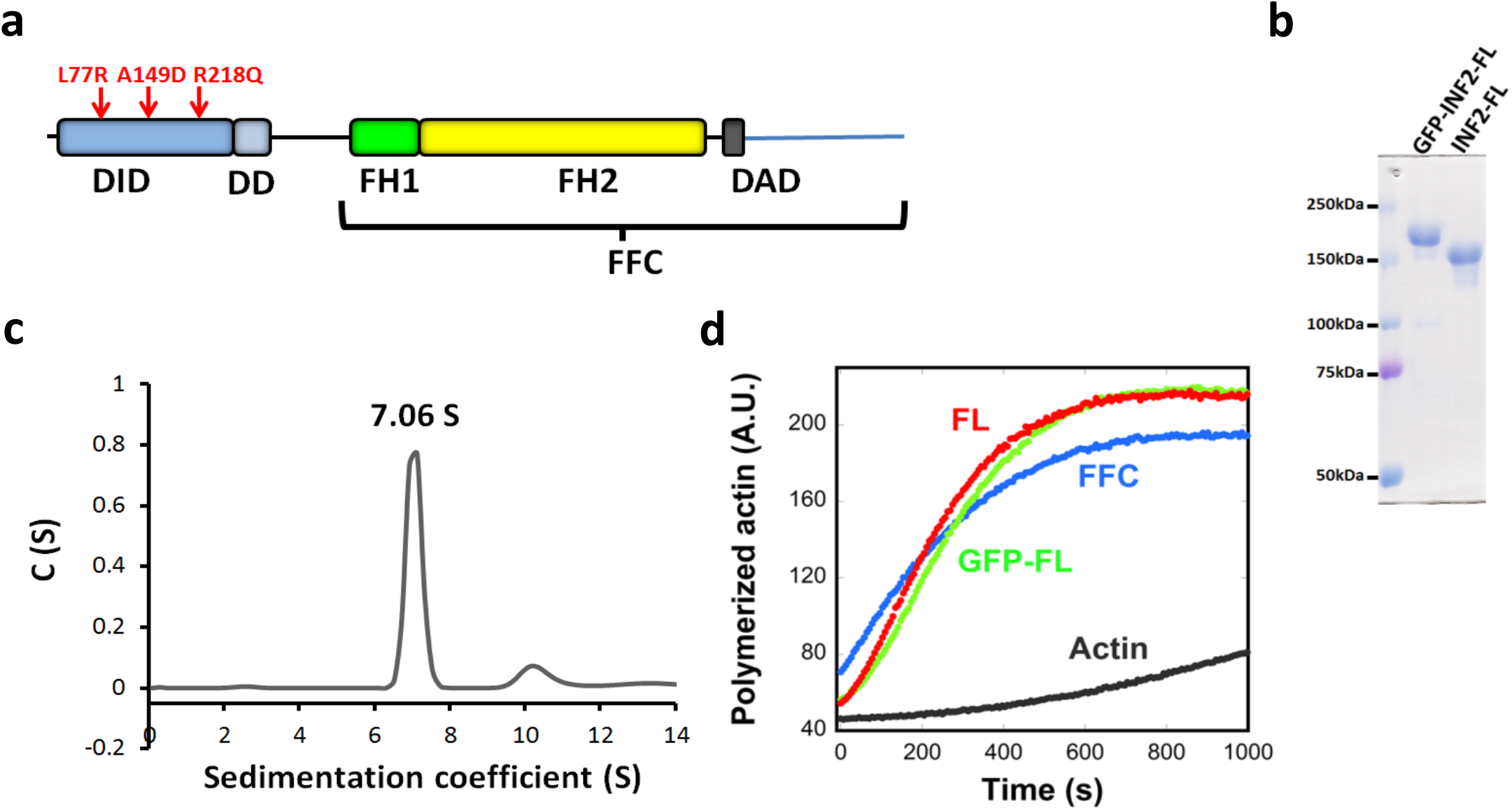
Purified INF2 full-length is constitutively active. **(a)** Schematic diagram of human INF2-nonCAAX (1240 residues). The FFC construct boundaries are 469-1240. The A149D mutation disrupts DID/DAD interaction. The R218Q and L77R link to FGSG and CMTD, respectively. We define domain boundaries as: DID, 32-261; DD, 263-342; FH1, 421 −520; FH2, 554-940; DAD, 971 −1000. **(b)** Coomassie-stained SDS-PAGE of human INF2-nonCAAX purified from HEK293 cells, with and without the GFP-Strep tag. **(c)** Sedimentation velocity analytical ultracentrifugation of INF2-nonCAAX (3.8 μM). Mass: 270.3 kDa (from sedimentation), 134.6 kDa (from sequence). **(d)** Pyrene actin polymerization assay (2*μ*M actin monomer, 5% pyrene label) using 20nM INF2 (with or without GFP-Strep) or INF2 FFC.

We also tested the effect of GFP-INF2-nonCAAX on cytosolic actin polymerization in U2OS cells, analyzing apical focal planes to avoid abundant ventral stress fibers (Fig. S1a). Wild-type INF2 causes no apparent increase in cytoplasmic actin filaments, while expression of a DID mutant predicted to disrupt DID/DAD interaction (INF2-A149D) increases cytoplasmic actin filaments significantly (Fig. S1b,c). These results agree with previous non-quantitative findings^30^, and suggest that INF2 is inhibited through DID/DAD interaction in cells, but that this interaction is insufficient for inhibition of the purified protein. One explanation could be that a cellular factor facilitates DID/DAD interaction. In this work, we identify such a factor.

### Identification of an INF2 inhibitory protein

Two possible mechanisms by which an inhibitor could enhance DID/DAD interaction are: (1) by direct binding; or (2) by post-translational modification (Fig. 2a). We used an activity-based biochemical approach to identify either mechanism, in which we mixed chromatographic fractions from mouse brain with purified INF2-nonCAAX, then assayed actin polymerization rate. We made two initial assumptions: 1) the inhibitor would be cytosolic, because both INF2-CAAX and INF2-nonCAAX are inhibited in cells; and 2) the inhibitor would be abundant, because even cells strongly over-expressing wild-type INF2 do not display aberrant actin polymerization (Fig. S1b, c).

**Figure 2.**
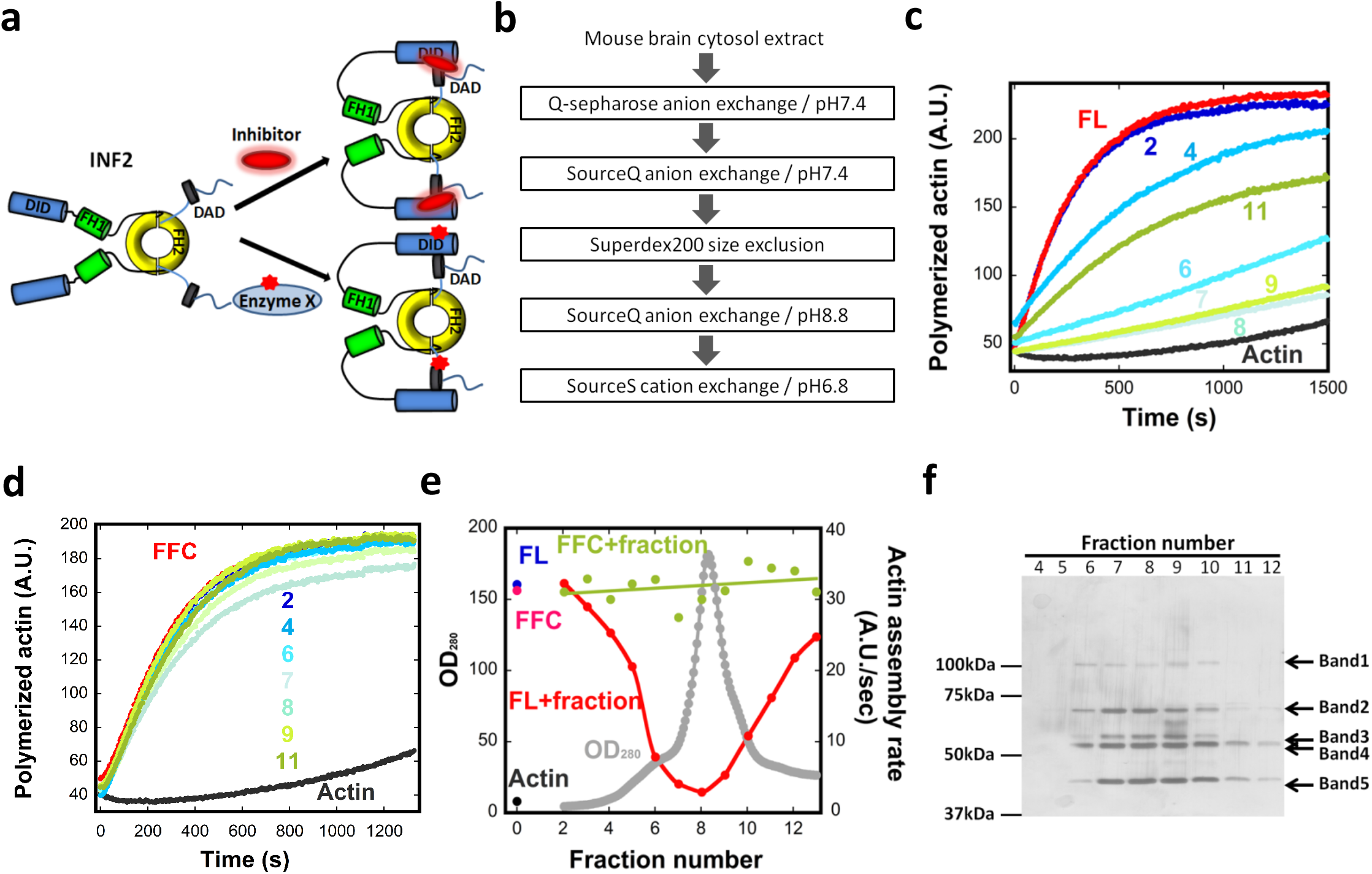
Identification of an INF2 inhibitor from mouse brain. **(a)** Schematic diagram of two possible mechanisms for facilitated autoinhibition of INF2. Upper: direct binding of inhibitor to INF2. Lower: post-translational modification of INF2 (red star). **(b)** Flow chart of column chromatography steps used for inhibitor purification. **(c)** Representative pyrene-actin assays containing 2μM actin monomer (5% pyrene) and 20nM INF2 full-length (FL) with or without the indicated BIF fractions (fraction #2-#11, from Fig. 2e). **(d)** Representative pyrene-actin assays containing 2μM actin monomer (5% pyrene) and 20nM INF2 FFC with or without the indicated BIF fractions (fraction #2-#11, from Fig. 2e). **(e)** Graph showing protein elution profile from the final SourceS column (OD280 nm, gray) as well as the activities of 20 nM INF2-FL (red) or FFC (green) in pyrene-actin assays on addition of the eluted fraction. Activity of INF2-FL alone (blue), INF2 FFC alone (magenta) and actin alone (black) are points on left side of graph. **(f)** Silver stained SDS-PAGE of indicated fractions from figure 2e. Bands identified as follows: Band1, Exportin1/7; Band 2, VATA and HSP7C; Band 3, CAP1, CAP2, VATB2; Band 4, CAP1, CAP2, VATA; and Band 5, actin.

Mouse brain cytosolic extract was fractionated through sequential column chromatography steps (Fig. 2b), pooling active fractions based on inhibition of INF2-FL activity in pyrene-actin assays (Fig. 2c,d, Fig. S2a-c). In the last chromatographic step, inhibitory activity eluted with the major protein peak (Fig. 2e). We refer to this fraction as Brain Inhibitory Fraction (BIF). Extensive optimization of the order and execution of the chromatographic steps was required to obtain the final BIF fraction, with the first SourceQ column being particularly important. Importantly, BIF does not inhibit actin polymerization by INF2-FFC (Fig. 2d,e) or mDia1-FFC (Fig. S2d), demonstrating that it is not acting as a general actin polymerization inhibitor. The inability of the BIF to inhibit INF2-FFC also demonstrates that DID is required for inhibition. BIF does not cause significant INF2 proteolysis (Fig. S2e), suggesting that the inhibitor is not a protease. BIF loses all activity upon boiling (Fig. S2f). These experiments reveal a protein inhibitor of INF2 that requires INF2’s DID for inhibition.

### The CAP/actin complex is an INF2 inhibitor

BIF contains five visible bands by SDS-PAGE and silver staining (Fig. 2f), which were identified by tryptic digest/mass spectrometry to be: Band 1, Exportin1/7; Band 2, Heat shock cognate 71kDa protein (HSP7C) and Vacuolar ATPase subunit A (VATA); Band 3, cyclase associated proteins 1 and 2 (CAP1, CAP2) and Vacuolar ATPase subunit B2 (VATB2); Band 4, CAP2, CAP1 and VATA; and Band 5, actin (Supplementary Table 1). We immuno-depleted individual proteins from BIF and tested the ability of these depleted fractions to inhibit INF2. Depletion of CAP2 eliminates INF2 inhibition (Fig. 3a,b), whereas depletion of VATA, VATB2 or HSP7C has no effect on inhibition (Fig. S3). Available antibodies against CAP1 and exportin proved unsuitable for immuno-depletion.

**Figure 3.**
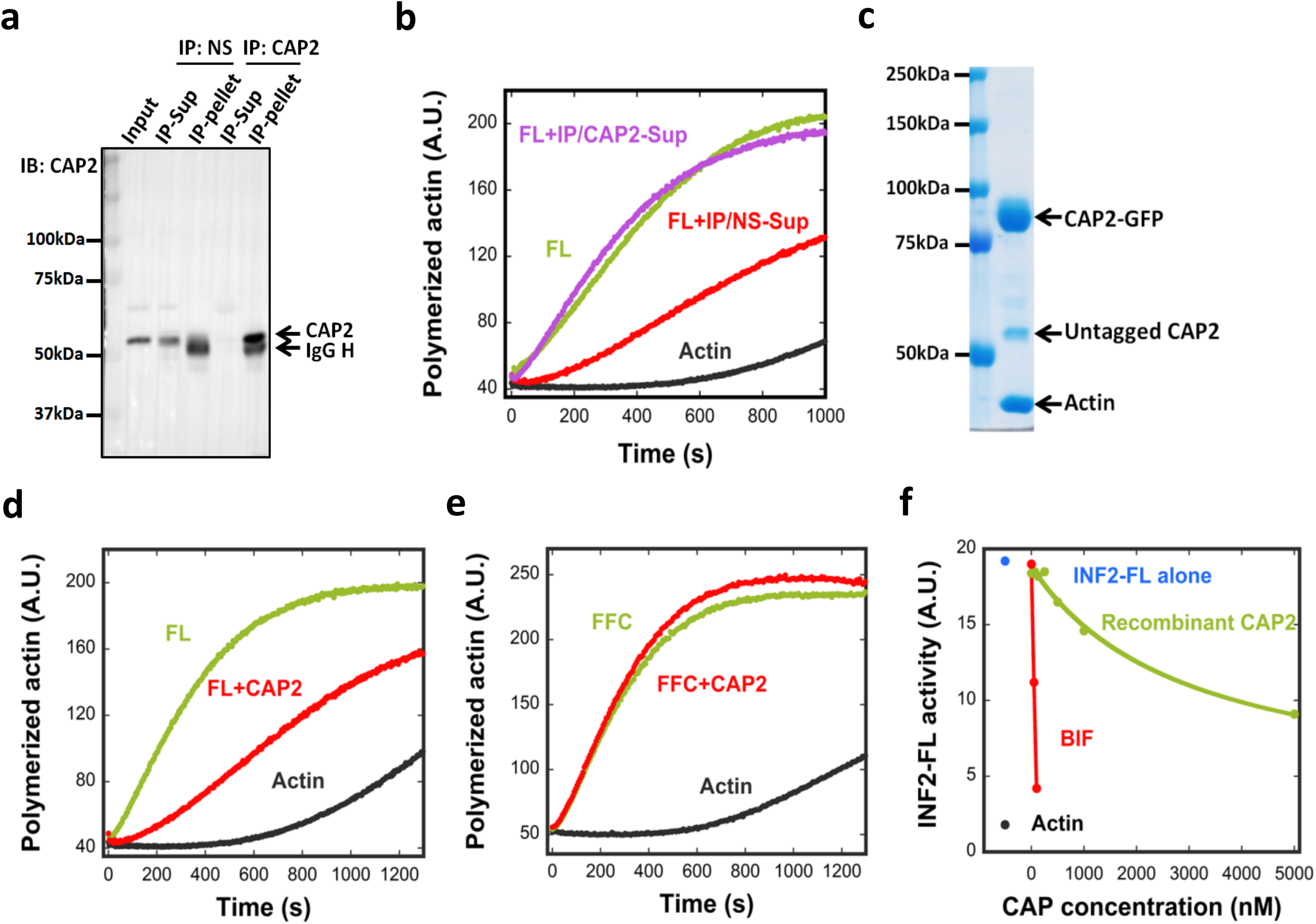
CAP/actin is an INF2 inhibitor. **(a)** Immuno-depletion of CAP2. An INF2-inhibiting fraction from mouse brain (Fraction 38-40 pool from SourceQ#1 (Fig. S2a)) was incubated with either a nonspecific AB (NS) or anti-CAP2 AB followed by precipitation using Protein A beads. CAP2 protein in input, supernatant and pellet detected by western blot. AB heavy chain, IgG H. **(b)** Pyrene-actin polymerization assay (2μM actin monomer, 5% pyrene) testing inhibitory activity of immune-depleted supernatants on 20nM INF2-FL. IP/NS-Sup: non-specific antibody supernatant. IP/CAP2-Sup: anti-CAP2 supernatant. **(c)** Coomassie stained SDS-PAGE of co-expressed CAP1-GFP/CAP2-GFP purified from HEK293 cells. **(d)** and **(e)** Pyrene-actin polymerization assay (2*μ*M actin monomer, 5% pyrene) testing effect of co-expressed CAP1/2 (5 μM, untagged) on 20nM INF2-FL **(d)** or on 20nM INF2 FFC **(e)**. **(f)** Comparing inhibitory activities of recombinant CAP (green) and CAP in BIF (red) on INF2-FL (20nM) in pyrene-actin assays. Assembly rate of actin alone (black) and with INF2-FL (blue) also shown. CAP concentration in BIF determined as described in Methods.

These results suggest that CAP2 is required for the INF2 inhibitory activity of BIF. Since additional peaks of INF2 inhibitory activity were identified in the first two purification steps (Fig. S2a,b), we used western blotting to ask whether CAP1 or CAP2 were present in these fractions. Both CAP1 and CAP2 were detected in inhibitory fractions from anion exchange chromatography (Fig. S2a) and size exclusion chromatography (Fig. S2b). These results suggest that endogenous CAP proteins are heterogeneous in biochemical properties, but that all CAP-containing peaks identified here can inhibit INF2.

To further test CAP as an INF2 inhibitor, we expressed and purified recombinant human CAP, containing C-terminal GFP and Strep-tag II (CAP-GFP), in HEK293 cells. We purified three versions of CAP: CAP1 alone, CAP2 alone, and co-expressed CAP1/CAP2. When analyzed by SDS-PAGE, all purified CAP-GFP fusions contain two additional bands: untagged CAP and actin (Fig. 3c, Fig. S4a). The mean molar ratio of untagged CAP to CAP-GFP is 0.16 in all cases, whereas the mean actin:CAP-GFP ratio is 0.88 (Table 1). Quantification of the protein bands in BIF reveals an actin:CAP ratio of ~1:1 (Table 1). CAP elutes as a high mass particle by gel filtration (Fig. S4b), and CAP-GFP sediments as a 14.5 S particle by analytical ultracentrifugation, with a calculated mass of 726 kDa (Fig. S4c). Considering past studies showing hexamerization of the N-termini of yeast and human CAP^33, 34^, we interpret our purified CAP-GFP to be a hexamer with a 5:1 ratio of CAP-GFP to untagged CAP, and actin monomer bound to a mean of five CAP subunits. The untagged CAP is likely endogenous CAP from HEK293 cells that has been incorporated into the CAP hexamer.

**Table 1.** Molar ratios in the CAP/actin complex. Calculated from densitometry of Coomassie-stained SDS-PAGE. CAP1, CAP2, and CAP1&2 = purified GFP-fusion proteins from HEK293 cells. BIF = purified brain inhibitory fraction. CAP-GFP (for CAP1, CAP2, and CAP1&2) or CAP (BIF) set to 100%.

| Mole percentage of indicated protein compared to recombinant CAP-GFP |  | CAP1 | CAP2 | CAP1&2 | BIF |
| --- | --- | --- | --- | --- | --- |
| Actin | Avg | 82.5% | 88.1% | 90.5% | 106.8% |
|  | STD | 9.4% | 11.7% | 10.7% | 4.3% |
|  | N | 6 | 24 | 2 | 3 |
| Untagged CAP | Avg | 16.8% | 16.3% | 16.5% | NA |
|  | STD | 14.3% | 12.4% | 15.0% | NA |
|  | N | 2 | 20 | 3 | NA |

Recombinant CAP/actin inhibits actin polymerization by full-length INF2 (Fig. 3d) but not by INF2-FFC (Fig. 3e, Fig. S4f). However, the inhibitory potency of recombinant CAP/actin is significantly lower than that of BIF (Fig. 3f). The IC_50_ for CAP in BIF is 69 nM, whereas recombinant CAP does not reach its IC_50_ at the maximum concentration tested. We obtain similar results for recombinant CAP1/actin and CAP2/actin, both with and without the GFP tag (Fig. S4d). In the absence of INF2, CAP/actin has little effect on polymerization of actin alone at the highest concentration tested (Fig. S4e). These results suggest that CAP is a component of the INF2 inhibitory factor, although recombinant CAP does not possess maximal inhibitory activity.

### Actin is an important component of CAP-mediated INF2 inhibition

We reasoned that the difference in inhibitory activity between recombinant CAP and BIF might be due to differences in the bound actin. To test this hypothesis, we attempted to exchange the bound actin in our recombinant CAP/actin (heretofore called CAP/293A) with other types of actin. We first tested our ability to exchange actin on CAP by mixing immobilized CAP/293A with fluorescently-labeled actin (TMR-actin) and monitoring the bead-bound fluorescence (Fig. S5a). Appreciable fluorescent actin associates with the CAP/293A-bound beads but not with control GFP-alone beads (Fig. S5b). SDS-PAGE analysis reveals that the ratio of actin:CAP is not changed significantly compared with mock-exchanged CAP beads (Fig. S5b). These results show that actin monomers can exchange onto CAP.

We exchanged CAP with total brain actin (BA), purified by a novel method (Fig. S5c). CAP/BA displays significantly more INF2 inhibition (IC_50_ 54 nM) than does CAP/293A (Fig. 4a,b), but does not inhibit INF2-FFC (Fig. 4c). We next exchanged CAP with two types of skeletal muscle actin, from chicken (CSKA) or from rabbit (RSKA). Interestingly, CAP/CSKA inhibits INF2-FL with an IC_50_ approaching that of CAP/BA (254 nM Fig. 4d,e), but does not inhibit INF2-FFC (Fig. S4f). In contrast, CAP/RSKA is a poor INF2 inhibitor, similar to CAP/293A (Fig. 4d,e). These results show that the source of the bound actin significantly affects INF2 inhibition by CAP.

**Figure 4.**
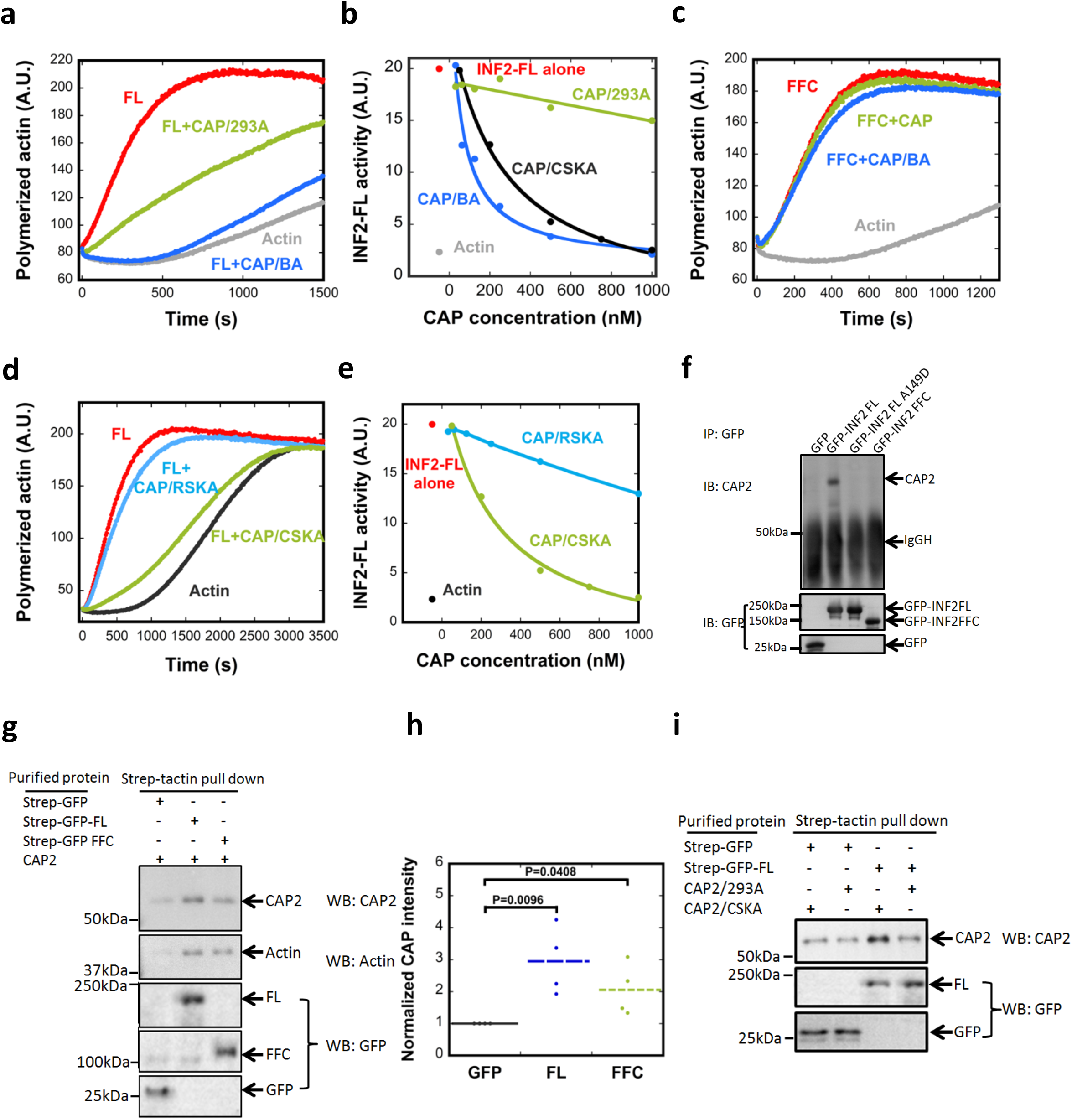
Actin-dependent differences in INF2 inhibition by CAP/actin. **(a)** Pyrene-actin polymerization assay (2*μ*M actin monomer, 5% pyrene) containing 20nM GFP-INF2-FL in the presence or absence of 5*μ*M purified CAP2-GFP (CAP/293A) or 250nM CAP2-GFP exchanged with brain actin (CAP/BA). **(b)** Concentration dependence of INF2 activity inhibition by CAP/293A, CAP/BA, or by CAP exchanged with chicken muscle actin (CAP/CSKA). **(c)** Pyrene-actin polymerization assay (2μM actin monomer, 5% pyrene) containing 20nM GFP-INF2 FFC in the presence or absence of 250nM CAP/293A or CAP/BA. **(d)** Pyrene-actin polymerization assay (2*μ*M actin monomer, 5% pyrene) containing 20nM GFP-INF2 full-length in the presence or absence of 1μM CAP/CSKA or CAP/RSKA (rabbit muscle actin). **(e)** Concentration dependence of INF2 inhibition by recombinant CAP2 exchanged with CAP/CSKA or CAP/RSKA. **(f)** Co-immunoprecipitation assay of endogenous CAP2 with transfected GFP-fusion proteins: GFP alone, GFP-INF2-FL, GFP-INF2FL A149D mutant, and GFP-INF2-FFC. Top panel: anti-CAP2 western on precipitated samples. Bottom panel: anti-GFP western on same samples. **(g)** Strep-tactin pull-downs performed with 1μM bead-bound strep-tagged GFP (control), strep-tagged GFP-INF2 full-length, or strep-tagged GFP-INF2 FFC. These were mixed with 2μM untagged CAP exchanged with chicken muscle actin (CAP/CSKA). Bound samples were analyzed by immunoblotting with anti-CAP2, anti-actin and anti-GFP. **(h)** Quantification of bound CAP2 from pull-downs as in Fig. 4g. CAP2 intensity from immunoblots normalized to strep-GFP pulldown. Bars, mean. Error bars, S.E.M. (n = 4 experiments). P values, one-sided student’s t-test. **(i)** Strep-tactin pull-downs performed with 1μM bead-bound strep-tagged GFP (control), or strep-tagged GFP-INF2 full-length. These were mixed with 2μM untagged CAP exchanged with chicken muscle actin (CAP/CSKA) or w/o exchange (CAP/293A). Bound samples analyzed by immunoblotting with anti-CAP2 and anti-GFP.

We next asked whether INF2 binds CAP/actin directly. First, we used a cellular co-IP approach where we transfected U2OS cells with GFP-INF2, then conducted anti-GFP IP and probed for CAP. CAP2 coprecipitates with GFP-INF2-FL but not with GFP-INF2-FFC nor with the A149D mutant of INF2-FL (Fig. 4f). This result suggests that DID/DAD interaction strongly enhances INF2 binding to CAP/actin. We also analyzed INF2/CAP interaction using purified proteins, by incubating immobilized GFP-INF2-FL or GFP-INF2-FFC with CAP/CSKA in solution, then analyzing the bead-associated CAP. CAP associates with INF2-FL to a greater extent than INF2-FFC (Fig. 4g,h). These results suggest that CAP/actin inhibits INF2 by direct interaction. In addition, CAP/CSKA binds INF2 better than does CAP/293A (Fig. S4i), consistent with their difference in inhibitory potency. These results suggest that the INF2/CAP interaction requires a specific type of actin.

Profilin is an abundant cytosolic actin-binding protein, and a significant proportion of cytosolic actin monomer is profilin-bound^35^. In addition, profilin binds CAP^36^. We tested the effect of profilin on INF2 inhibition by CAP/CSKA. By pyrene-actin polymerization, CAP/CSKA still potently inhibits INF2 in the presence of profilin (Fig. S5d), even though profilin slows the overall actin polymerization rate significantly. We also conducted total internal reflection (TIRF) microscopy assays of INF2-mediated actin polymerization in the presence of profilin, distinguishing between effects on actin nucleation and elongation. INF2 significantly increases nucleation rate, as judged by the number of actin filaments assembled over time (Fig. 5a,b), and increases the filament elongation rate approximately 2-fold (Fig. 5c). CAP/CSKA eliminates the INF2 effect on nucleation, while not having a significant effect on the nucleation rate of profilin/actin alone (Fig. 5a,b). CAP/CSKA has little effect on the elongation rate of profilin/actin alone, but inhibits the elongation rate of profilin/actin in the presence of INF2 (Fig. 5c). These results show that INF2 inhibition by CAP/actin can occur in the presence or absence of profilin.

**Figure 5.**
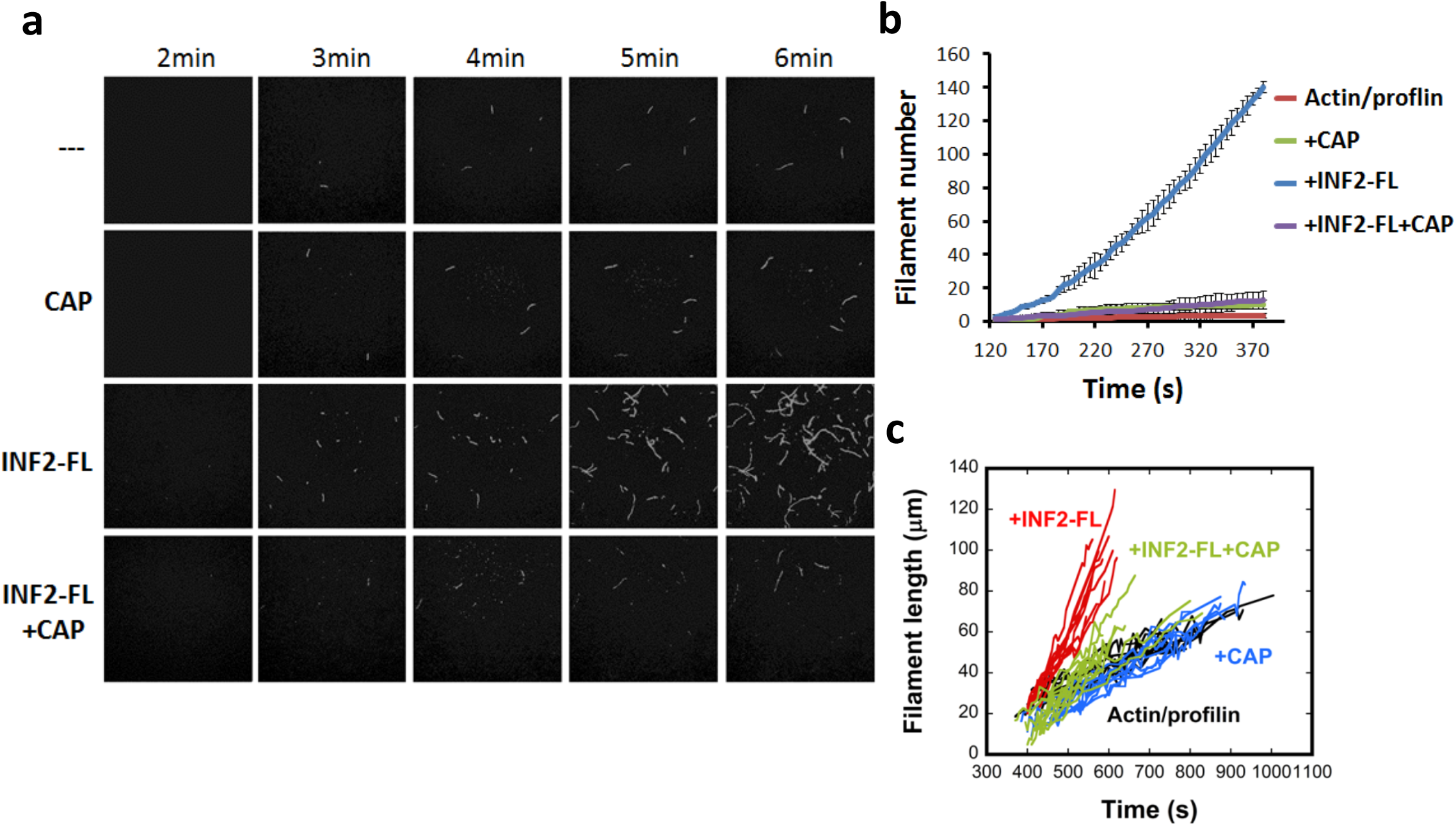
CAP inhibits INF2 mediated nucleation observed by TIRF microscopy. **(a)** Time-lapse TIRF microscopy images showing actin polymerization (1*μ*M actin monomer, 10% TAMRA-actin) containing 3*μ*M profilin in the presence or absence of 1nM INF2-FL and 50nM CAP2 exchanged with CSKA. **(b)** Quantification of filament number over time for TIRF assays (n=3 for INF2 FL, and n=5 for other conditions). **(c)** Measurement of filament length over time for TIRF assays. 10 filaments for each condition.

### Actin acetylation increases CAP/actin inhibition potency for INF2

We next probed the mechanism behind the differing potency between CAP/CSKA and CAP/RSKA for INF2 inhibition. Actin is highly conserved, with CSKA and RSKA being 100% identical in amino acid sequence. One possible explanation for the difference in INF2 inhibition is differential actin post-translational modification. By mass spectroscopy analysis, we identified acetylation on several lysines of actin for CAP/CSKA: K50, K61, K326 and K328 (Supplementary Table 2). We therefore tested whether lysine acetylation of actin was required for INF2 inhibition.

First, we attempted to deacetylate CAP/CSKA enzymatically. HDAC6 is a lysine deacetylase that resides in both the nucleus and cytoplasm^37^. We expressed and purified FLAG-tagged HDAC6 (Fig. S5e), which contains substantial deacetylase activity on purified brain tubulin (Fig. S5f). In pyrene-actin assays, HDAC6 strongly reduces CAP/CSKA inhibition of INF2, while minimally affecting polymerization of actin alone or in the presence of INF2 (Fig. 6a, FL+CAP+HDAC6 curve). To test whether HDAC6 deacetylase activity is required for this effect, we pre-treated HDAC6 with trichostatin A (TSA), an HDAC inhibitor. TSA pre-treatment abolishes the effect of HDAC6 on CAP/CSKA (Fig. 6a, FL+CAP+HDAC6/TSA curve). The concentration curves of CAP/CSKA treated with HDAC6 alone versus HDAC6 in the presence of TSA show a dramatic difference in their inhibitory activities (Fig. 6b). This result suggests that lysine acetylation on actin is necessary for CAP/actin inhibition of INF2.

**Figure 6.**
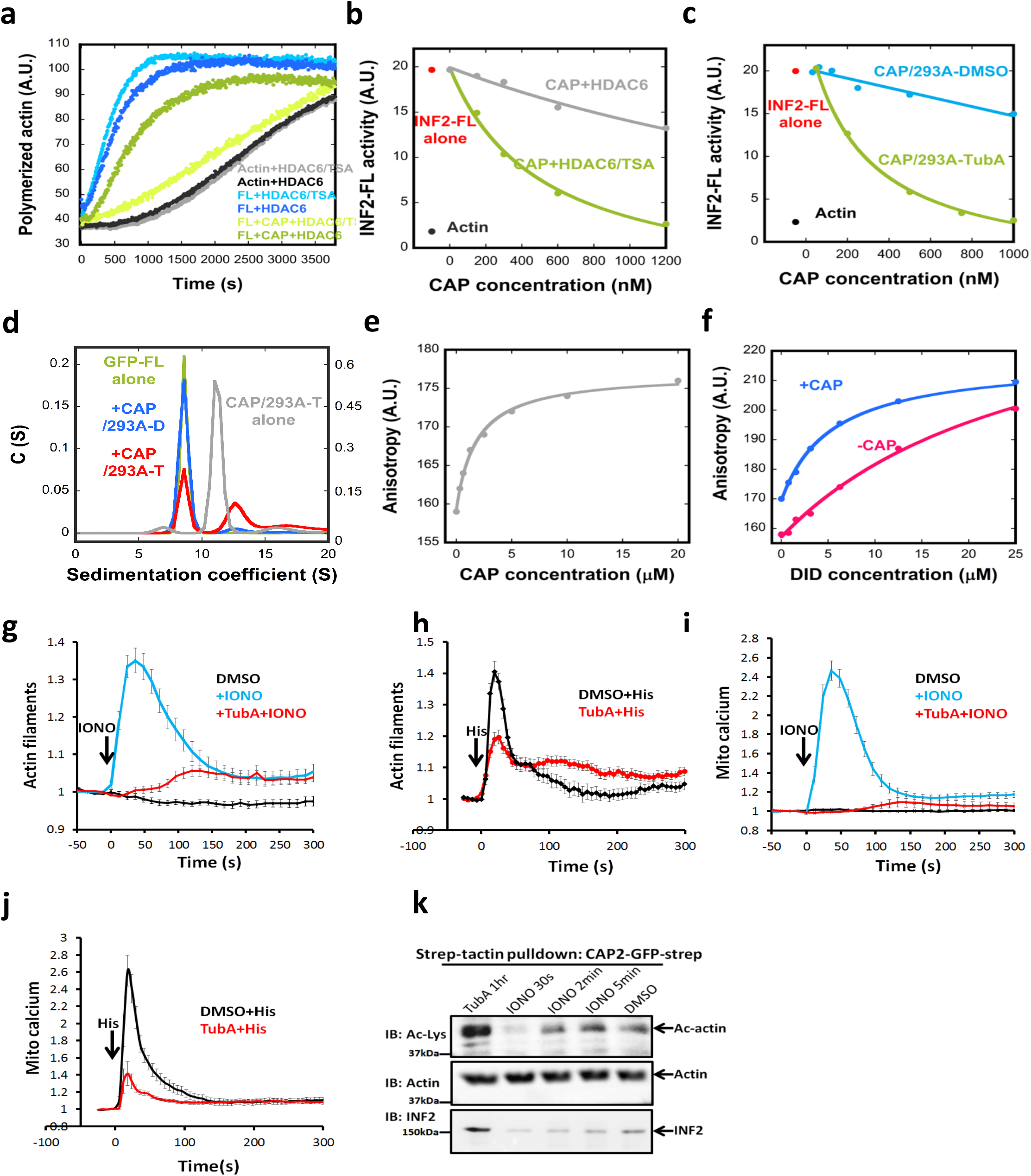
Acetylation-induced regulation of INF2 biochemically and in cells. **(a)** Pyrene actin polymerization assay (2μM actin monomer, 5% pyrene) containing 20nM GFP-INF2-FL in the presence or absence of 1μM CSKA-exchanged CAP2 (CAP2/CSKA) after pre-incubation with either HDAC6/TSA (HDAC6 with its inhibitor, TSA) or HDAC6 alone (active HDAC6). **(b)** Concentration dependence of CAP2/CSKA inhibition of INF2-FL after pre-treatment with HDAC6/TSA or HDAC6 alone. **(c)** Concentration dependence of INF2 inhibition by CAP2/293A purified from cells pretreated with DMSO or 50μM Tubastatin A for 1hr prior to lysis and purification. **(d)** Sedimentation velocity analytical ultracentrifugation of GFP-INF2-FL (1.8*μ*M), with or w/o 18*μ*M CAP2/293A-D or CAP2/293A-T. Mass of GFP-INF2-FL: 325.2 kDa (from sedimentation), 161.3 kDa (from sequence). **(e)** Fluorescence anisotropy assay for CAP2/293A-T binding by INF2-Cterm (100 nM). **(f)** Fluorescence anisotropy assay for INF2-Nterm binding to INF2-Cterm (100 nM) in presence or absence of CAP2/293-T (20 μM). Additional point at 48 μM INF2-Nterm is not shown, but is accounted for in the fit curve for both conditions. **(g)** Ionomycin-induced changes in actin filament levels in U2OS cells pretreated with either DMSO or 50*μ*M TubA. 4*μ*M ionomycin. Error bars, S.E.M. N for DMSO, 40; IONO, 38; IONO+TubA, 48. **(h)** Histamine-induced changes on actin filaments in U2OS cells pretreated with either DMSO or 50*μ*M TubA. 100*μ*M histamine. Error bars, S.E.M. N for Histamine, 15; Histamine+TubA, 15. **(i)** Ionomycin-induced changes on mitochondrial calcium levels in U2OS cells pretreated with either DMSO or 50*μ*M TubA. 4*μ*M ionomycin. Error bars, S.E.M. N for DMSO, 29; IONO, 28; IONO+TubA, 22. **(j)** Histamine-induced changes on mitochondrial calcium in U2OS cells pretreated with either DMSO or 50*μ*M TubA. 100*μ*M histamine. Error bars, S.E.M. N for Histamine, 15; Histamine+TubA, 15. **(k)** Strep-tactin pull-downs performed with cell lysates of U2OS cells overexpressing CAP2-GFP-2xStrep. Cells were treated with indicated conditions prior to lysis. CAP2-GFP was pulled down and CAP2-bound actin and its acetylation level were analyzed by immunoblotting with anti-actin and anti-Ac-Lys antibodies.

We took a second approach to test HDAC6’s effect on CAP/actin, by purifying CAP/actin from 293 cells in which HDAC6 activity is inhibited. Cells were treated either with DMSO or with the HDAC6-specific inhibitor tubastatin A (TubA) for 1hr before cell harvesting and purification. Two tests indicate the effectiveness of TubA treatment. First, the amount of acetylated tubulin in cell extracts increases dramatically, as monitored by western blotting with anti-acetyl-K40-tubulin (Fig. S5g). Second, the amount of acetylated lysine detected on actin in the purified CAP/293A complex increases, as monitored by western blotting with a pan-anti-acetyl-lysine antibody (Fig. S5h). By pyrene-actin assay, the TubA-treated CAP/293A (CAP/293A-T) inhibits INF2 with an IC_50_ of 263nM, which is significantly more potent that the DMSO-treated sample (Fig. 6c). These results suggest that 1) HDAC6 de-acetylates actin in cells, and 2) increased actin acetylation increases the inhibitory potency of CAP/actin.

We used the high inhibitory potency of CAP/293A-T, versus the mock-treated CAP/293A-D (DMSO), to analyze the direct interaction between INF2 and CAP/actin in more detail. By analytical ultracentrifugation, the presence of CAP/293A-T causes a portion of GFP-INF2-FL to shift to a higher S species (Fig. 6d, red curve). The sedimentation coefficient of the larger species, 12.6 S, is greater than that of the un-tagged CAP/293A-T complex (11.2 S, gray curve), suggesting INF2 binding to the hexameric CAP complex. CAP/293A-D does not produce this magnitude of shift in GFP-INF2 (Fig. 6d, blue curve), suggesting its lower affinity. In addition, no shift in S value occurs for GFP-INF2-FFC in the presence of CAP/293A-T (Fig. S5i), suggesting that DID is required for high-affinity interaction.

We also used fluorescence polarization of fluorescently-labeled INF2-Cterm (containing the DAD) to examine its binding to INF2-Nterm (containing the DID) in the presence or absence of CAP/293A-T. INF2-Cterm displays some affinity for CAP/293A-T, with a Kd^app^ of 1.82 μM (Fig. 6e). Intriguingly, the affinity of INF2-Cterm for INF2-Nterm increases at least 5-fold in the presence of CAP/293A-T (Fig. 6f), with the Kd^app^ of DID/DAD estimated at 28 μM and 5.4 μM in the absence and presence of CAP/293A-T, respectively (see Methods for explanation of Kd^app^ determination). These results suggest that the CAP/actin complex increases the affinity between DID and DAD.

We used mass spectroscopy to compare acetylation patterns on actin for the CAP/293A-T and the CAP/293A samples. Using relative peak areas, four acetylation sites on actin are appreciably enriched in the TubA-treated samples (K50, K68, K215, and K315), while three sites display no clear change in acetylation (K191, K326, K328, Supplementary Table 3). These results suggest that the increase in CAP/actin inhibitory potency might be linked to acetylation of one or more of these residues (K50, K68, K215 and K315).

The ability of HDAC6 to abolish CAP/actin inhibition of INF2 suggests that HDAC6 might be required for cellular INF2 activation. Previously, we and others have shown that cell stimulation with ionomycin or histamine causes an INF2-dependent cytosolic actin “burst”^23, 38-40^. We tested the effect of TubA pretreatment on the actin burst in U2OS cells. TubA pre-treatment strongly reduces the actin burst caused by ionomycin (Fig. 6g) or histamine (Fig. 6h). A down-stream consequence of INF2 activation is increased mitochondrial calcium, leading to inner mitochondrial membrane constriction and mitochondrial division^23^. TubA pre-treatment strongly reduces the mitochondrial calcium increase caused by ionomycin (Fig. 6i) or histamine (Fig. 6j). These results show that HDAC6 can activate INF2 both biochemically and in cells.

We also tested whether ionomycin treatment alters lysine acetylation of actin in cells. Since western blotting of Ac-K residues from whole-cell lysates resulted in high background in the region of actin, we took an alternate approach in which we expressed 2xStrep-tagged CAP2 in U2OS cells, then affinity-purified the CAP2 and examined the Ac-K levels on the bound actin by western blot at various time points after treatment. Mock treatment with DMSO results in a measurable level of Ac-K, and treatment with TubA strongly increases Ac-K level (Fig. 6k). Ionomycin treatment causes a significant drop in Ac-K at 30 sec, with a return to control levels at 2 min (Fig. 6k). This time course of Ac-K decrease correlates with the time course of the ionomycin-induced actin burst (Fig. 6g). In addition, endogenous INF2 co-purifies with CAP2 in the DMSO-treated and the TubA-treated samples, but co-purification is greatly reduced at 30 sec post-ionomycin treatment (Fig. 6k). These results suggest that ionomycin causes a transient decrease in lysine acetylation of actin, resulting in decrease INF2 binding.

### INF2 disease mutants have reduced sensitivity to CAP/actin inhibition

Although multiple mutations in INF2’s DID have been linked to two human diseases, FSGS and CMTD, the molecular mechanism of disease progression is unknown. We therefore asked whether INF2 disease mutants were inhibited by CAP/KAc-actin complex. For this analysis, we chose two DID mutants: R218Q, the most common FSGS mutant^28, 41, 42^; and L77R, a CMTD mutant^29^. R218 is located near the DAD-binding site but is not predicted to make direct contact with DAD^28^, while L77 is localized on the opposite face of the DID from the DAD binding site. At 1 μM, CAP/CSKA displays reduced inhibition of L77R-INF2 (27% decrease versus >90%) and no measurable inhibition of R218Q-INF2 (Fig. 7a). A concentration curve with CAP/293A-T shows potent inhibition of WT-INF2 but not of either mutant, with R218Q being particularly resistant (Fig. 7b). These results show that INF2 disease mutants are not efficiently regulated by CAP/KAc-actin.

**Figure 7.**
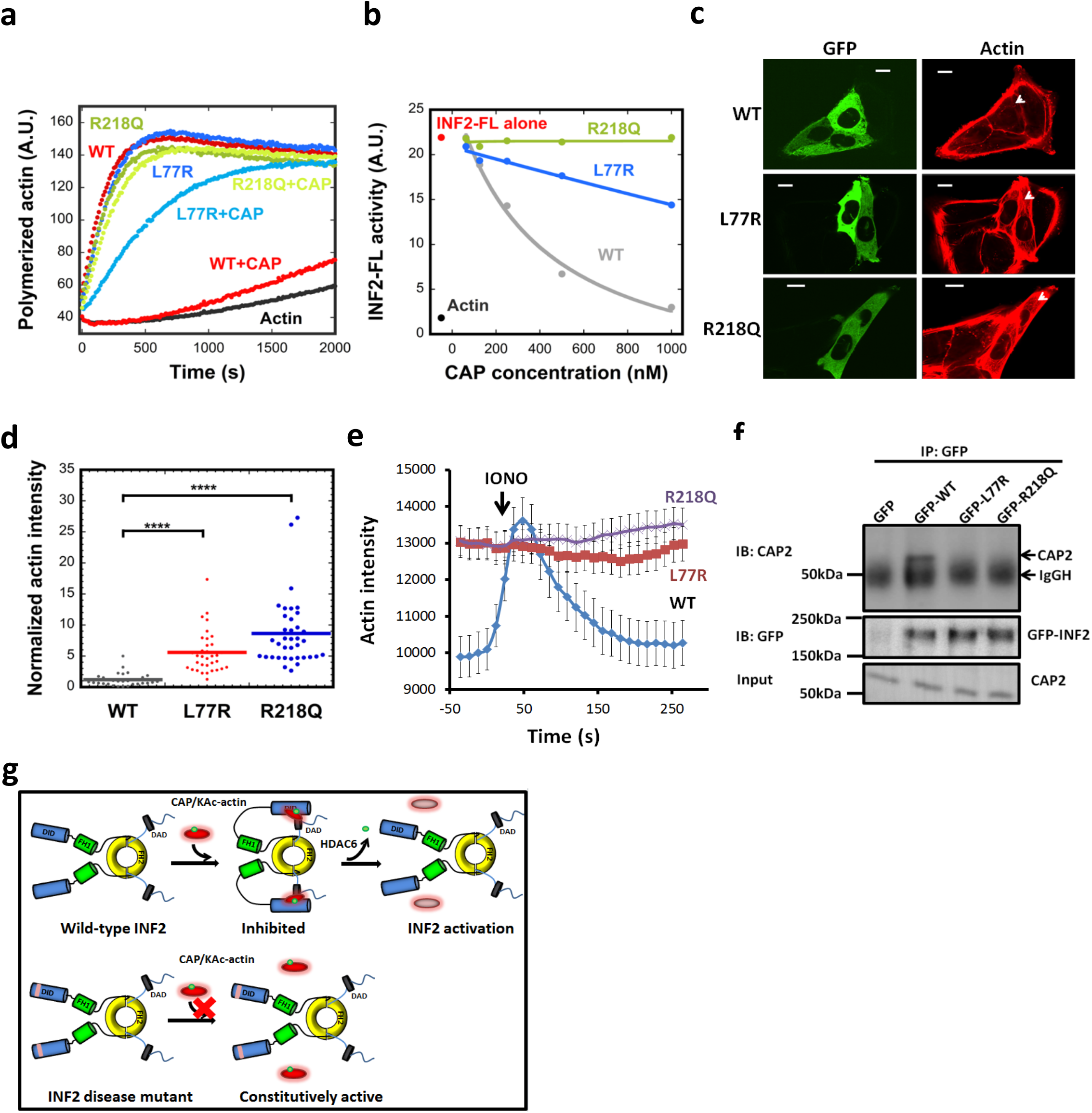
INF2 disease mutants display decreased regulation by CAP/actin. **(a)** Pyrene-actin polymerization assay (2μM actin monomer, 5% pyrene) containing 20nM GFP-INF2 wildtype (WT) or mutants (FSGS mutant R218Q, CMTD mutant L77R) in presence or absence of 1*μ*M CAP2/CSKA. **(b)** Concentration dependence of CAP2/293A-T inhibition of INF2-FL and disease mutants, in pyrene-actin assays conducted as in panel a. **(c)** Representative GFP and TRITC-phalloidin confocal microscopy images showing cytosolic actin polymerization by exogenously expressed GFP-INF2 constructs (WT, L77R, or R218Q) in INF2-KO U2OS cells. Cells were fixed and stained with TRITC-phalloidin to label actin filaments. Images are maximum intensity projections of three confocal imaging planes in the middle z region of the cell. Bar, 10 *μ*m. **(d)** Normalized fluorescence intensity of TRITC-phalloidin was quantified from images similar to **(c)**. Lines represent mean ***(WT: n=31 cells, mean=1.19, STD=1.02; L77R: n=34 cells, mean=5.60, STD=3.40; R218Q: n=37 cells, mean=8.63, STD=5.47)*.** *^****^ denotes p* < 0.0001 by one-sided student’s t-test. **(e)** Time course of ionomycin-induced changes in actin filaments in INF2-KO U2OS cells expressing GFP-fusions of INF2 WT or the indicated mutants along with mApple-Ftractin. Error bars, S.E.M. N = 34, 35, and 33 cells for WT, L77R and R218Q, respectively. **(f)** Co-immunoprecipitation assay of endogenous CAP2 with transfected GFP-fusion proteins: GFP alone, GFP-INF2-FL, GFP-INF2FL L77R mutant, and GFP-INF2FL R218Q mutant. Top panel: anti-CAP2 western on precipitated samples (IgGH, IgG heavy chain from the IP). Middle panel: anti-GFP western showing GFP-INF2 in precipitated samples. Bottom panel: anti-CAP2 western of input samples. **(g)** Schematic model of INF2 facilitated autoinhibition by CAP/KAc-actin, and activation by HDAC6-mediated actin de-acetylation. INF2 mutants linked with FSGS and CMTD are defective in CAP/KAc-actin binding, leading to constitutive activity.

We also tested the regulation of L77R- and R218Q-INF2 in cells, by transfecting the mutant nonCAAX constructs into INF2-null U2OS cells. Fixed-cell analysis shows that both mutants display significantly higher cytosolic actin filaments than WT-INF2 (Fig. 7c,d). Live-cell analysis shows that cytoplasmic actin filament levels are significantly higher for both mutants than for WT-INF2, and that ionomycin stimulation does not induce an increase in actin filaments for the mutants (Fig. 7e). We also tested the interaction between these INF2 mutants and CAP2 in cells, and found no apparent co-immunoprecipitation, in contrast to INF2-WT (Fig. 7f). Finally, we tested the effect of TubA treatment on cytoplasmic actin filament levels in L77R-INF2 or R218Q-INF2 expressing cells, and found no decrease in these levels, suggesting that HDAC6 inhibition, which should increase lysine acetylation of actin, does not inhibit these mutants efficiently (Fig. S5j,k). These results show that two INF2 disease mutants are constitutively active for actin polymerization in cells.

## DISCUSSION

We reveal the first cellular role for lysine-acetylated actin: inhibition of INF2 while in complex with CAP. This mechanism is novel for the regulation of actin dynamics in cells, and we refer to the mechanism as ‘facilitated autoinhibition’ because an additional molecule is required to mediate INF2’s autoinhibitory interaction. Actin de-acetylation through HDAC6 relieves CAP/actin-mediated INF2 inhibition. Cellular HDAC6 inhibition blocks stimulus-induced INF2 activation. Conversely, INF2 mutants linked to two human diseases, FSGS and CMTD, are poorly inhibited by CAP/KAc-actin and are constitutively active in cells.

### Roles for lysine acetylation in cytoskeletal regulation

A variety of actin post-translational modifications have previously been identified, with methionine oxidation and arginylation being the best described to-date^43^. Actin acetylation research has mainly focused on the N-terminus, where all actins are modified^44^. Lysine acetylation on actin has been identified^43^ but its significance has been unclear until this study.

Our demonstration that lysine-acetylated actin inhibits INF2 adds to known acetylation events for other cytoskeletal proteins. Two actin-binding proteins have been shown to be acetylated: cortactin^45^, and the formin mDia2^46^. In both cases, acetylation inhibits the interaction of these proteins with actin, decreasing actin polymerization. For microtubules, acetylation of lysine-40 on a-tubulin has effects on microtubule structure^47^, while β-tubulin acetylation might inhibit microtubule polymerization^48^. Interestingly, formin proteins can influence tubulin acetylation through a variety of mechanisms^20, 49, 50^.

### Cyclase-associated protein as a component of the INF2 inhibitory complex

The presence of CAP in the INF2 inhibitory complex is intriguing. CAP was originally identified in budding yeast as an interacting protein for adenylyl cyclase^51, 52^, but this interaction is not conserved in mammals. Subsequently, CAP was shown to bind actin monomers and filaments^53-56^. Purified CAP stimulates actin filament disassembly in the presence of cofilin or twinfilin, but CAP alone has little effect on actin disassembly^34, 57^. The two mammalian CAPs, CAP1 and CAP2, are ~60% identical and are differentially expressed^58, 59^. Both CAP1 and CAP2 inhibit INF2. Three results indicate that CAP’s role in INF2 regulation is directly on INF2, as opposed to effects on actin: purified CAP/actin has little effect on the polymerization of actin alone (Fig. S4e, Fig. 5), CAP/actin does not inhibit INF2 that is lacking the DID (Fig. 4c), and CAP/actin associates with INF2 (Fig. 4f-i, Fig. 6d-f).

CAP/actin is a high molecular weight complex (Fig. S4b,c), consistent with previous reports of hexamerization through the CAP N-terminus^33, 34^. Interestingly, additional fractions from mouse brain that inhibit INF2 also contain CAP1 or CAP2 (Fig. S2a,b). The gel filtration profiles of these fractions (Fig. S2b) suggest that cellular CAP is heterogeneous, and might include some non-hexameric species. A second possibility is that CAP complexes with additional proteins. CAP is currently known to interact with several actin-binding proteins, including cofilin, profilin, twinfilin and ABP1^34, 57, 60^. The fraction we purified as BIF contained several co-eluting proteins, including subunits of vacuolar ATPase and exportin. It is possible that these putative binding proteins might provide additional properties.

### Facilitated autoinhibition of INF2 by the CAP/KAc-actin complex

Auto-inhibition through DID/DAD interaction is a major predicted regulatory mechanism for most metazoan formins^4-6^. In addition, some formins not containing canonical DID or DAD are still subject to direct auto-inhibition through N-terminal/C-terminal interaction^61^. Finally, a number of proteins can inhibit formins in trans, binding at or near the FH2 domain^62-68^.

Our results demonstrate the first example of facilitation of formin auto-inhibition by another protein factor. The fact that a mutation known to disrupt DID/DAD interaction results in constitutive INF2 activity (Fig. S1) suggests that DID/DAD interaction is required. However, we cannot rule out the possibility that CAP/actin serves as a bridge between DID and DAD, with DAD binding KAc-actin directly, due to its high affinity for actin^15, 30^. This interaction would require release of CAP’s WH2 from KAc-actin, and would also block canonical DID/DAD interaction. The Kd^app^ of the DAD-containing INF2-Cterm for CAP/actin (1.82 μM) is 10-fold higher than that for rabbit muscle actin alone (0.18 μM^30^), and would have to displace CAP’s WH2 motif from the actin. The fact that CAP can associate with INF2 (Fig. 4g-i) suggests that CAP is part of the inhibitory complex and not merely delivering KAc-actin. Our schematic model of an inhibitory complex between CAP/actin and INF2 (Fig. 7g) does not distinguish between direct DID/DAD interaction or the CAP/actin bridge model, nor does it depict CAP hexamerization, two issues that must be addressed in future work.

The acetylation positions identified on actin provide insight into the inhibitory mechanism of the CAP/KAc-actin complex. Four lysines display appreciable acetylation increases in the inhibitory CAP/actin complex: K50, K68, K215 and K315. These residues are all on the surface of the actin monomer (Fig. S6a), although K315 likely contributes to actin monomer stability through interactions between subdomains. These residues do not make significant inter-subunit contacts in the filament (Fig. S6b), so acetylation may not disrupt polymerization. Similarly, acetylation is not predicted to alter CAP binding affinity, since none of the four lysines make direct contact with the actin-binding CARP domain of CAP (Fig. S6c), or the predicted binding site of CAP’s WH2 motif^69^. Current models of FH2/actin structure also suggest little interaction of these residues with the FH2 (Fig. S6d).

### Cellular regulation of INF2 through actin acetylation

Several questions arise as to cellular INF2 regulation through this mechanism. First, what percentage of actin must be acetylated for INF2 inhibition? We have previously determined total cytosolic actin to be ~200 μM in U2OS cells^70^, consistent with values in other cell types^35^, whereas INF2 levels are <100 nM (unpublished results). Therefore, acetylation of <0.1% of cytosolic actin could be sufficient for INF2 inhibition.

A second question concerns the cellular signal that triggers actin deacetylation and INF2 activation. It is clear that an increase in cytoplasmic calcium activates INF2^23, 38-40^, but the mechanism linking increased cytoplasmic calcium to decreased actin acetylation is unclear at present. One possibility is through calcium-mediated changes in metabolism^71^. Regulation of acetylation is governed by the balance between lysine acetyltransferase (KAT) and lysine deacetylase activities, and the appreciable cellular acetylation/deacetylation flux^37^ could be due either to a decrease in KAT activity or an increase in deacetylase activity. Several cytosolic members of the ~20 mammalian KAT protein family could mediate actin acetylation^37^. An intriguing possibility is that fluctuating levels of the acetyl donor, acetyl-CoA, could regulate KAT activity^72^. Such a mechanism links well with INF2’s role in mitochondrial division^23, 25^ and the relationship between mitochondrial division and metabolism^73^. Alternately, increased calcium might increase deacetylase activity. Our evidence using HDAC6 inhibitors in cells suggest that HDAC6 is the relevant actin deacetylase for INF2 activation. Interestingly, HDAC6 activity has been shown to cause an increase cellular actin polymerization^45, 74^. Another possibility is that calmodulin might participate in calcium-mediated INF2 activation. An interaction between INF2 and calcium-bound calmodulin has been shown in cell lysates^40^. However, we have not detected calmodulin in our preparations of CAP/KAc-actin complex or INF2, and we do not find an effect of purified calmodulin on INF2 activity (not shown).

### INF2 regulation and human disease

Our results have exciting implications for disease mechanisms of both FSGS and CMTD. All disease-linked INF2 mutations map to the DID, but few are at the DAD-binding site, suggesting that they do not affect this interaction directly. In addition, all disease mutations are dominant, and expression of select mutations in cells causes aberrant actin polymerization (Fig. 7c,^28^), suggesting that both FSGS and CMTD might be due to increased INF2 activity. We show that two mutants, one for FSGS and one for CMTD, are poorly inhibited by CAP/KAc-actin. Therefore, mutations that weaken INF2 binding to the CAP/KAc-actin inhibitory complex may lead to disease through increased INF2 activity. Interestingly, recent studies have shown that aberrant HDAC6 activity can also lead to CMTD^75-77^, raising the possibility of another route for INF2 activation in this disease. Overall, our study provides a framework for understanding both actin regulation and disease progression in a new way.

## METHODS

### Plasmids

Human INF2 full-length nonCAAX cDNA was made from the human INF2-CAAX clone (Origene SC313010, NM_022489) followed by replacement of the CAAX exon with the nonCAAX exon, correction of a 12-base deletion in the purchased clone, and codon optimization. This construct was subsequently cloned into the EcoR1/Xho1 sites of a modified eGFP-C1 vector (Clonetech Inc) containing Strep-tag II (IBA Life Sciences) and an HRV3C cleavage site N-terminal to the INF2 start codon. INF2 mutations (A149D, L77R, R218Q) were generated in the above-mentioned INF2 full-length nonCAAX construct by site-directed mutagenesis. Human CAP1 and CAP2 cDNAs were purchased from NovoPro (710829-5 (NM-006367) and 710470-11 (NM-006366)), then the entire ORF was PCR-amplified and cloned into a modified eGFP-N1 vector (Clonetech) containing an HRV3C cleavage site followed by the GFP ORF, followed by Strep-tag II, using Xho1 and Kpn1 sites. FLAG tagged human HDAC6 plasmid was purchased from Addgene (30482). Mito-R-GECO1, Cyto-R-GECO1 (K_d_= 0.48 *μ*M for calcium) constructs were kind gifts from Yuriy M. Usachev (Dept of Pharmacology, University of Iowa Carver College of Medicine), are on the pcDNA3.1 (-) backbone (Thermo Fisher), and have been described previously ^78^. GFP-F-tractin and mApple-F-tractin plasmids are described in ^79, 80^.

### Recombinant protein expression and purification

For protein expression in mamalian cells, 2.24 x 10^6^ FreeStyle 293-F cells (Life Tech R790-07) growing in 1L FreeStyle 293 expression medium (Life Tech 12338-018) were transfected with 0.5mg DNA and 1.5mg sterile 25kDa linear PEI mixed in Opti-MEM reduced-serum medium (Life Tech 31985070). Proteins were expressed at 37 °C, 8% CO2 with shaking at 125 rpm for 2 days (INF2) or 3 days (CAP1, CAP2 and HDAC6). For the Tubstatin A treatment of CAP2-transfected cells, cells were centrifuged at 300xg at 4°C after 3 days of transfection, and resuspended in Dulbecco modified Eagle media (DMEM) without serum. DMSO or 50 μM Tubastatin A (Selleckchem S2627) was added and cells were cultured for an additional 1hr before harvest.

For strep-tagged protein purification from FreeStyle 293 cells, all following steps were performed at 4°C or on ice. Cells were pelleted at 300xg for 15min, and pellet was resuspended in 45mL EB (100mM Hepes pH7.4, 500mM NaCl_2_, 5mM EDTA, 1mM DTT, 1% v/v TritonX-100, 2μg/mL leupeptin, 10μg/mL aprotinin, 2μg/mL pepstatin A, 1μg/mL calpeptin, 1μg/mL calpain inhibitor 1, 1mM bezamidine, 1:1000 dilution of universal nuclease) per 5mL cell pellet followed by 30min end-over-end incubation. For tubastatin A-treated cells, EB also contained 500 nM tricostatin A ***(Selleck S1045)*** and 10 mM sodium butyrate ***(Sigma-Aldrich 303410)*.** The cell debris was removed by ultracentrifugation at 185,000xg for 1 hr (Ti45 rotor, Beckman), and the supernatant was blocked with avidin (20μg/mL Sigma-Aldrich 189725) then applied to Strep-Tactin Superflow resin (2-1206-025; IBA, Göttingen, Germany) column equilibrated in EB by gravity flow. After thorough wash with WB (10mM Hepes pH7.4, 150mM NaCl2, 1mM EDTA, 1mM DTT), column was either: 1) treated with HRV3C protease in WB (at 1:50 enzyme to substrate molar ratio) for 16hrs at 4°C followed by 3 column volume WB wash to obtain untagged protein; or 2) eluted with strep elution buffer (10mM Hepes pH7.4, 150mM NaCl, 1mM EDTA, 1mM DTT, 2.5mM desthiobiotin) to obtain strep-GFP fusion protein. Protein was concentrated with 30,000 molecular-weight-cutoff Amicon Ultra-15 filter unit (Millipore) before further purification through Superdex 200 16/60 gel-filtration column (GE Bioscience) with final buffer (10mM Hepes pH7.4, 50mM KCl, 1 mM MgCl_2_, 1mM EGTA, 1 mM DTT). Protein was further concentrated to >10 μM before being aliquoted, frozen in liquid nitrogen, and stored at −80°C. N.B. – in all cases, Hepes pH given is the pH at 23°C.

For FLAG-tagged protein purification from Freestyle 293 cells, cells were harvested the same way as described above, and lysed in EB-F (50 mM Hepes 7.4, 150 mM NaCl, 1 mM EDTA, 1 mM DTT, 1% v/v TritonX-100, 2μg/mL leupeptin, 10μg/mL aprotinin, 2μg/mL pepstatin A, 1μg/mL calpeptin, 1μg/mL calpain inhibitor 1, 1mM bezamidine, 1:1000 dilution of universal nuclease). The cell lysate was clarified by ultracentrifugation at 185,000xg for 1 hr (Ti45 rotor, Beckman), and supernatant was loaded onto Anti-FLAG M2 affinity gel equilibrated with EB-F. The column was washed with WB-F (10 mM Hepes 7.4, 50 mM KCl, 1mM MgCl_2_, 1 mM EGTA, 1 mM DTT) thoroughly, and FLAG-tagged protein was eluted with FLAG elution buffer (10 mM Hepes 7.4, 50 mM KCl, 1mM MgCl_2_, 1 mM EGTA, 1 mM DTT, 100μg/mL FLAG octapeptide (Sigma F3290)). Protein was gel filtered, concentrated and stored as described above.

### Actin preparation

Rabbit or chicken skeletal muscle actin was purified from acetone powder ^81^. Rabbit muscle actin was labeled with pyrenyliodoacetamide ^82^, TAMRA NHS ester (Molecular probes; C1171) or Oregon488 NHS ester (Thermo Fisher; O6185) ^83^. Both labeled and unlabeled actin were gel-filtered on Superdex 75 (GE Biosciences) and stored in G-buffer (2 mM Tris-HCl pH 8, 0.5 mM DTT, 0.2 mM ATP, 0.1 mM CaCl_2_, 0.01% w/v sodium azide) at 4°C.

For brain actin purification, all steps were at 4°C. 50g mouse brains were homogenized using polytron (Kinematica PT3100) in EB (50mM Hepes pH7.4, 1mM MgCl_2_, 1mM EGTA, 2mM DTT, 2μg/mL leupeptin, 10μg/mL aprotinin, 2μg/mL pepstatin A, 1μg/mL calpeptin, 1μg/mL calpain inhibitor 1, 1mM bezamidine). The cell debris was removed by ultracentrifugation at 185,000xg (Ti45 rotor), and the supernatant was filtered through 0.45μm filters. Supernatant was supplemented with ATP (4mM), MgCl_2_ (4mM), and sodium phosphate (NaPO_4_, pH 7.0, 20 mM), followed by end-over-end incubation at 4°C for 16hrs. Polymerized brain actin was pelleted by centrifugation for 3hrs at 185,000xg, and the pellet was washed by resuspension-pelleting cycle sequentially with 2xWB1 (500mM KCl, 0.5% v/v thesit, 10mM NaPO_4_ pH7.0, 2mM MgCl_2_, 2mM ATP, 1mM DTT, 2μg/mL leupeptin), and 2xWB2 (50mM KCl, 10mM NaPO_4_ pH7.0, 2mM MgCl_2_, 2mM ATP, 1 mM DTT, 2 μg/mL leupeptin). Pellet was then washed with G-buffer with minimal disturbance and then resuspended in G-buffer before shearing through 30G 1/2 needle. Sheared actin was dialyzed in G-buffer for 16hrs before ultracentrifugation 185,000xg (Ti70.1 rotor). After the spin, supernatant was concentrated to >10μM.

### BIF purification

BIF purification was conducted at 4°C. 100g mouse brains were homogenized using polytron (Kinematica PT3100) in EB (50mM Hepes pH7.4, 1mM MgCl_2_, 5mM EGTA, 1mM DTT, 2μg/mL leupeptin, 10μg/mL aprotinin, 2μg/mL pepstatin A, 1μg/mL calpeptin, 1μg/mL calpain inhibitor 1, 1mM bezamidine). The cell debris was removed by ultracentrifugation at 185,000xg (Ti45 rotor), and the supernatant was filtered through 0.45μm filters. The supernatant was supplement with KCl to 100mM before loading onto 90mL Q-sepharose column (GE Biosciences), BIF containing fraction was eluted with Q350 (350mM KCl, 50mM Hepes pH7.4, 1mM MgCl_2_, 5mM EGTA, 1mM DTT, 2μg/mL leupeptin, 10μg/mL aprototin, 2μg/mL pepstatin A, 1μg/mL calpeptin, 1 μg/mL calpain inhibitor 1). Eluate was diluted 3-fold with EB before loading onto 50mL Source Q column (GE Biosciences). Linear ionic strength gradient elution was performed with Q100 (100mM KCl, 10mM Hepes pH7.4, 1mM MgCl_2_, 1mM EGTA, 1mM DTT) and Q350 (350mM KCl, 10mM Hepes pH7.4, 1mM MgCl_2_, 1mM EGTA, 1mM DTT). Fractions were dialyzed into 1.5xKMEH+DTT (15mM Hepes pH7.4, 75mM KCl, 1.5mM MgCl_2_, 1.5mM EGTA, 1mM DTT) and tested for inhibition of INF2 full-length activity by pyrene actin polymerization assay (20 nM INF2, 2 μM actin (5% pyrene)). Inhibitory peak fractions were pooled and purified through Superdex 200 16/60 gel filtration column (GE Biosciences) equilibrated in 1.5xKMEH+DTT. Inhibitory activity was assayed again and inhibitory peak fractions were pooled and purified through 8mL Source Q column at pH8.8 with gradient from Q100 (100mM KCl, 10mM Tris pH8.8, 1mM MgCl_2_, 1mM EGTA, 1mM DTT) to Q300 (300mM KCl, 10mM Tris pH8.8, 1mM MgCl_2_, 1mM EGTA, 1mM DTT). Fractions were dialyzed into 1.5xKMEH_DTT and inhibitory activity was assayed. Peak inhibitory fractions were pooled and dialyzed into Q0 (10mM Hepes pH6.8 at 4°C, 1mM MgCl_2_, 1mM EGTA, 1mM DTT) before applied to 1mL Source S column (GE Biosciences) at pH6.8 with gradient elution from Q0 to Q100 (10mM Hepes pH6.8 at 4C, 100mM KCl, 1mM MgCl_2_, 1mM EGTA, 1mM DTT). Fractions were dialyzed into 1.5xKMEH+DTT, assayed for inhibitory activity, analyzed by silver-stained SDS-PAGE and pooled.

For CAP concentration determination in BIF, 75 μL sample from 1mL Source S column fractions (Fig. 2E) was mixed with 25uL SDS-PAGE sample buffer and resolved along with actin standards with known mass by large 7.5% SDS-PAGE. Gel was stained with Colloidal Blue Staining SDS-PAGE, and CAP concentration was analyzed by densitometry (ImageJ), using a standard curve of known actin concentrations. For determination of IC_50_ for CAP in BIF, linear fit was applied to activity of BIF in Fig. 3F, and activity of actin was used as baseline.

### TIRF Microscopy

Methods have been described in detail elsewhere^17^. Briefly, glass flow chambers were assembled using VWR micro coverglasses (22×22 and 18×18 mm, No 1.5) with double-stick Scotch tape to hold 10μL volume. Before assembly, cover glasses were washed in acetone (50 min), ethanol (10 min), and MilliQ water (1 min) and then incubated in Pirhana Solution (1:2 ratio of 30% H_2_O_2_ and H_2_SO_4_) for 1 h. Glasses were then rinsed with water, followed by 0.1M KOH, followed by water again and dried with inert gas before silanization. Glasses were silanized overnight in a 0.0025% solution of dichlorodimethyl silane (Sigma-Aldrich 85126) in chloroform, washed with methanol, dried with inert gas, and stored in clean sealed containers. Before flowing into samples, chambers were incubated with 1% w/v Pluronic F127 (Sigma-Aldrich P2443) in BRB80 (80mM PIPES/KOH, pH 6.9, 1 mM EGTA, 1 mM MgCl_2_) for 1 min and then equilibrated in TIRF buffer (10mM imidazole pH7, 50mM KCl, 1 mM MgCl_2_, 1 mM EGTA, 1 mM DTT, 100 mM DTT, 0.2 mM ATP, 15 mM glucose, 0.5% methylcellulose, 0.01mg/ml catalase [Sigma C3515], 0.05mg/ml glucose oxidase [Sigma G6125], and 0.1% BSA). Nucleation-elongation assays were conducted as previously mentioned^16, 17^. Briefly, unlabeled rabbit skeletal muscle actin monomers were mixed with 20% TAMRA or Oregon-labeled actin monomers (2μM) in G buffer, diluted with profilin (3μM) in 2xTIRF buffer in the presence or absence of 1 nM untagged INF2 full-length construct and/or 100nM CAP/CKSA, and introduced into the flow chamber. The filaments were visualized on a Nikon Eclipse Ti-E inverted microscope with 488- or 561-nm lasers and driven by Nikon Elements software. Single-color images were acquired every 5 s at 100 ms exposure with TIRF objective (60X1.49 N.A.) and Andor electron-multiplying charge-coupled device camera with an Andor TuCam adapter using Perfect Focus. Filament number and length were measured manually from frame to frame using Nikon NIS-Elements. For display of TIRF images, original images were adjusted using photoshop CS5. Curve function was used to lock the intensity of the filaments and then reduce the background haze. All the panels were adjusted as a single image.

### Pyrene actin polymerization assay

Rabbit skeletal muscle actin in G-buffer (6 μM actin, 5% pyrene) was converted to Mg^2+^ salt by addition of 1 mM EGTA and 0.1 mM MgCl_2_ for 2 min at 23°C immediately prior to polymerization. Polymerization was induced by addition of 2 volumes of formin with or without other proteins (INF2 and/or inhibitory fraction) in 1.5xpolymerization buffer (75mM KCl, 1.5mM MgCl_2_, 1.5mM EGTA, 15 mM Hepes pH 7.4, 2mM DTT, 2 mM Tris-HCl, 0.2 mM ATP, 0.1 mMCaCl_2_, and 0.01% w/v NaN_3_). Pyrene fluorescence (365/410 nm) was monitored in a 96-well fluorescence plate reader (Infinite M1000; Tecan, Mannedorf, Switzerland) within 1 min of mixing actin and formin or other proteins/fraction. Slope of curve at ½ max is plotted to represent INF2 activity.

### Velocity Analytical Ultracentrifugation

Analytical ultracentrifugation was conducted using a Beckman Proteomelab XL-A and an AN-60 rotor. For sedimentation velocity analytical ultracentrifugation, untagged INF2 full-length nonCAAX (3.8μM) or CAP2-GFP (7μM) in K50MEHD (10mM Hepes pH7.4, 50mM KCl, 1mM MgCl_2_, 1mM EGTA, 1mM DTT) was centrifuged at 25,000 rpm with monitoring at 490 nm. Data were analyzed by Sedfit to determine sedimentation coefficient, frictional ratio, and molecular weight. Sedimentation coefficient reported is that of the major peak (at least 80% of the total analyzed mass) at OD_280_ for untagged INF2 or CAP2 or OD_490_ for GFP-INF2 or CAP2-GFP.

### Antibodies

Vacuolar ATPase subunit A (rabbit anti-ATP6V1A, Abcam, ab199325), Vacuolar ATPase subunit B2 (rabbit anti-ATP6V1B2, Abcam, ab73404), heat shock cogate 71kDa protein (rabbit anti-Hsc70, Abcam, ab51052), CAP2 (rabbit polyclonal, Santa Cruz, sc-167378), CAP1 (rabbit polyclonal, Abcam ab96354), GFP (rabbit polyclonal, gift from William Wickner (Dartmouth)), actin (mouse monoclonal, clone C4, Millipore, MAB1501) alpha-tubulin (mouse monoclonal, clone DM1A, Sigma-Aldrich, T9026), acetyl-lysine (rabbit polyclonal, Cell Signaling Technologies, 9441), anti-acetylated tubulin (mouse monoclonal, clone 6-11B-1 against acetylated lysine 40 of α-tubulin, Sigma-Aldrich, T7451). Anti-INF2 was described previously (^27^; 941-1249 antibody used, rabbit polyclonal).

### Immuno-depletion

The initial SourceQ inhibitory fraction pool from mouse brain prep (fractions 38-40, Fig. S2A) was incubated with antibody against Vacuolar ATPase subunit A, Vacuolar ATPase subunit B2, heat shock cogate 71kDa protein, CAP2, or control IgG (rabbit anti-GFP), respectively at 4°C. 10μL of each antibody was added into 500μL fraction followed by 16hrs incubation with end-over-end rotation at 4°C. The antigen-antibody complex was precipitated by protein A or protein G agarose beads (GE). Supernatant was recovered and tested for INF2 inhibition. Supernatants and pellets were probed for the depleted protein.

### CAP bound actin exchange assay

CAP2-GFP-2xstrep protein purified from 293 cells (also containing bound actin from 293 cells) was immobilized on strep-tactin beads, which were then washed with G-buffer. The indicated species of actin (in G-buffer) were added and incubated with CAP2-GFP strep-tactin beads at 4°C with end-over-end rotation for 12hrs. Concentrations of added actin and bead-bound CAP2-GFP in exchange reactions were 5μM and 1 μM, respectively. Beads were washed with 20 column volumes of G-buffer. Exchanged CAP2-GFP was removed from beads by either: eluting with G-buffer containing 2.5mM dethiobiotin, or 2) cleavage from beads by HRV3C protease (1:50, overnight at 4°C).

### Protein binding assays

For strep-tactin pulldown of purified proteins, 1 μM strep-GFP tagged protein (GFP, INF2FL or INF2FFC) was incubated with 2μM untagged CAP (with or w/o actin exchange) in IPB (50mM KCl, 1 mM MgCl_2_, 1 mM EGTA, 10 mM Hepes pH 7.4, 1mM DTT, 1% v/v thesit) with end-over-end rotation at 4°C for 8hrs. Separately, 0.2 volumes of 50% strep-tactin bead slurry were blocked with 10mg/mL avidin and washed in PD before adding to protein mix. After 8hr incubation at 4°C, beads were washed with IPB (now containing 0.2% v/v thesit). The bead pellet was resolved by SDS-PAGE, and the western blot probed with anti-GFP, anti-CAP2, or anti-actin antibodies.

For strep-tactin pulldown from cell extracts, Wild-type U2OS cells expressing CAP2-GFP were treated with 50μM Tubstatin A for 1hr or with DMSO for 15min or with 4μM ionomycin (in serum-containing medium) for the indicated length of time prior to lysis. Cells were lysed immediately after removal of media with prechilled PDEB (50mM KCl, 1mM MgCl_2_, 5mM EGTA, 10 mM Hepes pH 7.4, 1mM DTT, 3% v/v thesit, 2μg/mL leupeptin, 10μg/mL aprototin, 2μg/mL pepstatin A, 1μg/mL calpeptin, 1μg/mL calpain inhibitor 1, 1mM bezamidine, 500nM Tricostatin A, 10mM sodium butyrate) in the dish and then incubated with shaking at 4°C for 15min. The lysate was spun at 100K rpm with TLA120 rotor using Beckman tabletop ultracentrifuge.at 4°C for 20min, and supernatant was blocked with avidin (20μg/mL Sigma-Aldrich 189725) then mixed with Strep-Tactin Superflow resin (2-1206-025; IBA, Göttingen, Germany) prewashed with EB with end-over-end rotation for 3hrs at 4C. Beads were washed with PDEB twice and 1xPBS twice. The pellet was resolved by SDS-PAGE, and the western blot was probed with anti-Actin, anti-acetylated lysine or anti-INF2 antibodies.

For immunoprecipitation of purified proteins, 0.5μM strep-GFP tagged protein (GFP, INF2-FL or INF2-FFC) was incubated with 2μM untagged CAP (with or w/o exchange) in IPB for 3hrs at 4°C. 50μg anti-GFP antibody was added and the mix was incubated 12hrs with end-over-end rotation at 4°C. Protein A beads (GE) were blocked with 10mg/mL avidin and washed in IPB before incubating with protein mix. After another 3hrs incubation, beads were washed with IPB (now containing 0.2% v/v thesit). The bead pellet was resolved by SDS-PAGE, and the western blot was probed with anti-GFP, anti-CAP2, or anti-actin antibodies. For statistical analysis, results from three independent experiments were quantified. CAP2 band intensity was normalized against GFP control using ImageJ.

For immuno-precipitation from cell extracts, INF2KO U2OS cells expressing GFP, GFP-INF2 full-length nonCAAX WT, GFP-INF2FFC or GFP-INF2 full-length nonCAAX A149D, GFP-INF2 full-length nonCAAX L77R, GFP-INF2 full-length nonCAAX R218Q mutants were trypsinzed, centrifuged at 300xg for 5 min, and the cell pellet washed with PBS. Cell pellet was lysed with prechilled IPEB (50mM KCl, 1mM MgCl_2_, 5mM EGTA, 10 mM Hepes pH 7.4, 1mM DTT, 3% v/v thesit, 2μg/mL leupeptin, 10μg/mL aprototin, 2μg/mL pepstatin A, 1 μg/mL calpeptin, 1 μg/mL calpain inhibitor 1, 1mM bezamidine), then treated with latrunculin A (Sigma-Aldrich, L5163) at 20μM for 15min on ice. Cell debris were removed by ultracentrifugation using TLA-100 (Beckman) at 100,000 rpm. Supernatant was incubated with 10μg anti-GFP with end-over-end rotation at 4°C for 12hrs. Protein A beads were pre-blocked with 10mg/mL avidin (sigma189725) and washed in IPEB before incubating with lysate for 3hr. 20 μ L Beads were washed with IPW (50mM KCl, 1mM MgCl_2_, 1mM EGTA, 10 mM Hepes pH 7.4, 1mM DTT, 0.2% v/v thesit) and bound proteins were resolved by SDS-PAGE western, probed with anti-GFP and anti-CAP2.

Fluorescence polarization measurements were conducted using INF2-Cterm (amino acids 941-1249 of INF2-CAAX, expressed in bacteria) labeled with tetramethylrhodamine-succinimide (TMR), as previously described^30^. TMR-INF2-Cterm (100 nM) was mixed with CAP/293A-T and/or INF2-Nterm (amino acids 1-420, expressed in bacteria as described^30^) in 50mM KCl, 1mM MgCl_2_, 5mM EGTA, 10 mM Hepes pH 7.4, 1mM DTT at 23°C, and fluorescence anisotropy measured in the Tecan M1000 plate reader. The concentration of CAP/293A-T was held constant at 20 μM in some assays. Binding curves were fit using standard hyperbolic saturation fitting^84^. In the case of the INF2-Cterm/INF2-Nterm interaction in the absence of CAP/293A-T, saturation was not obtained and had to be assumed based on extrapolation of the curve, which adds uncertainty to the K_d_^app^ value of 28 μM determined for this interaction. For this reason, we state that CAP/293A-T increases the affinity of INF2-Nterm for INF2-Cterm at least 5-fold, which is a conservative estimate of this increase.

### Mass spectrometry

Samples were separated by SDS-PAGE, bands were excised, destained, and digested with trypsin in 50 mM Ammonium bicarbonate overnight at 37°C. Peptides were extracted using 5% formic acid/50% ACN and dried. Peptides were analyzed on a Fusion Orbitrap mass spectrometer (ThermoScientific) equipped with an Easy-nLC 1000 (ThermoScientific). Raw data were searched using COMET in high resolution mode ^85^ trypsin enzyme specificity with up to three missed cleavages, and carbamidomethylcysteine as fixed modification. Oxidized methionine, and acetylated lysine were searched as variable modifications. Quantification of LC-MS/MS spectra was performed using MassChroQ ^86^. Peptides were corrected based on protein amount, ratios CAP TubA / DMSO were calculated on a per charge state and per peptide basis and averaged. Additional analyses were performed at the Taplin Mass Spectrometry Facility at Harvard.

### Live cell imaging

Wild-type human osteosarcoma U2OS cells (American Type Culture Collection HTB96) or INF2 KO U2OS cells (described in ^23^) were grown in DMEM (Invitrogen) supplemented with 10% calf serum (Atlanta Biologicals). Cells tested every 6 months for mycoplasma contamination using LookOut PCR detection kit (Sigma-Aldrich). Cells were seeded at 4×10^5^ cells per well in a 6-well dish ~16 hours prior to transfection. Plasmid transfections were performed in OPTI-MEM media (Invitrogen) with 2 μL Lipofectamine 2000 (Invitrogen) and 500 ng GFP-Ftractin and/or mito-R-GECO1, or GFP-INF2 full-length nonCAAX, GFP-INF2 full-length nonCAAX L77R mutant or GFP-INF2 full-length nonCAAX R218Q mutant with mApple-Ftractin per well for 6 hours, followed by trypsinization and re-plating onto glass bottom MatTek dishes (P35G-1.5-14-C) at ~2×10^5^ cells per well. Cells was imaged in live in DMEM (GIBCO, #21063-029 with 4.5g/L D-Glucose, L-Glutamine and 25Mm HEPES) supplemented with 10% NCS (HyClone, #SH30118.03) ~16-24 hours after transfection. For histamine or ionomycin treatments, cells were treated with 100 *μ*M histamine (Sigma Aldrich H7125, from 100 mM stock in DMSO), 4 *μ*M ionomycin (Sigma Aldrich I0634, from 2 mM stock in DMSO), or DMSO alone at 1 min after commencement of imaging, and imaging and continued for another 5-10 min. Medium was pre-equilibrated for temp and CO_2_ content before use. For Tubastatin A (Selleckchem S2627) treatments, cells were pretreated with either 50 *μ*M Tubastatin A (from 10 mM stock in DMSO) or equal volume DMSO (control) for 60 min prior to addition of histamine or ionomycin as above. Note: Tubastatin A-containing media was made immediately prior to addition to the cells, individually for each plate. Imaging was conducted on a Dragonfly 302 spinning disk confocal (Andor Technology Inc, Belfast UK) on a Nikon Ti-E base and equipped with an iXon Ultra 888 EMCCD camera, and a Tokai Hit stage-top incubator. Lasers: solid state 405 smart diode 100 mW, solid state 488 OPSL smart laser 50 mW, solid state 560 OPSL smart laser 50 mW, solid state 637 OPSL smart laser 140 mW. Objective: 100x 1.4 NA CFI Plan Apo (Nikon). Images acquired using Fusion software (Andor).

### Fix cell imaging

For fixed cells, cells were transfected with GFP, GFP-INF2 full-length nonCAAX, GFP-INF2 full-length nonCAAX A149D mutant, GFP-INF2 full-length nonCAAX L77R mutant or GFP-INF2 full-length nonCAAX R218Q mutant in the same way as for live cell, and split onto MatTek dishes. Cells were then treated with DMSO or 50 μM Tubastatin A for 1hr or without any treatment prior to fixation with 4% paraformaldehyde (Electron Microscopy Sciences Inc) in phosphate-buffered saline (PBS) for 20min at room temperature. After washing with PBS, the cells were permeabilized on ice with 0.25% Triton X-100 in PBS for 15min. Cells were then washed with PBS prior to blocking with 10% calf serum in PBS for 1hr at room temperature. Actin was stained with 100nM TRITC-phalloidin (Sigma-Aldrich) and DNA was stained with 0.1mg/L w/v DAPI (Calbiochem 268298) for 10min at room temperature in dark followed by PBS wash. Cells were imaged on the MatTek dishes in PBS using the same confocal microscope as for live-cell (Dragonfly) but with Zyla 4.2 Mpixel sCMOS camera, and z stacks with 0.2μm stepsize were taken from bottom of cells to the top. Images of from middle z planes were used for quantification. Cells were selected based on GFP expression, then imaged for both GFP and phalloidin staining without first examining the degree of phalloidin staining.

### Measurements of actin intensity in fixed cells

TRITC-phalloidin intensity of two ROIs in the perinuclear region were taken per cell (medial z-section taken), and was normalized by the intensity of 2 ROIs taken from nuclear region (which lacked measurable phalloidin staining). 30 cells for each condition were quantified.

### Measurements of calcium changes and actin burst in live cells

Measurements of the following parameters were conducted as described previously^23^. Briefly, cells were seeded at 4×10^5^ confluency and transfected 24 hours prior to imaging with the indicated mitochondrial calcium and Ftractin probes. Cells were treated with inhibitors and/or stimuli as described above. Imaging was conducted in a medial region of the cell, approximately 2 μm above the basal surface. Mean fluorescence was calculated for each cell using ImageJ (NIH). Fluorescence values for each time point after drug treatment (F) were normalized by the average initial fluorescence (first 5-6 frames prior to stimulation-F0) and plotted against time as F/F0.

#### Calcium measurements

All cells in the imaging field were analyzed. Where cells were closely juxtaposed, their calcium changes were analyzed in the same ROI. Numbers of cells quantified in histamine experiments: cytosolic calcium: histamine control 14; tubastatin A: 36, and mitochondrial calcium: histamine control: 15 cells; tubastatin A: 15 cells. Number of cells quantified in ionomycin experiments: cytosolic calcium: DMSO control 20; ionomycin control: 19; tubastatin A: 17, and mitochondrial calcium: DMSO control 29; ionomycin control: 28; tubastatin A: 22.

#### Actin “burst” measurements

Mean Fluorescence values for each time point were calculated from 4 ROI/cell selected in the peri-nuclear region using ImageJ. Only cells fully in the imaging field were analyzed. Cells were excluded from analysis for reasons: they were rounded up, or a significant concentration of basal stress fiber signal in the field contributed to abnormally high initial signal. Excluded cell percentages as follows. Figure 6d: 15%, 16% and 19% for DMSO, +Iono and +TubA+Iono, respectively. Figure 6e: 32% for both DMSO+histamine and TubA+histamine. Fluorescence values for each time point after drug treatment (F) were normalized with the average initial fluorescence (first 5-6 frames prior to treatment-F0) and plotted against time as F/F0. Numbers of cells quantified in histamine experiments: histamine control 18 cells; tubastatin A: 30 cells. Number of cells quantified in ionomycin experiments: DMSO control 40; ionomycin control: 38; tubastatin A: 48.

### Statistical Analysis

Error (standard error of the mean) was calculated using Excel (Microsoft, version 16.16.1). Comparisons by one-sided T-test (paired) were conducted using GraphPad.

## Data Availability

The mass spectrometry proteomics data (Supplementary Tables 1,2, and 3) have been deposited to the ProteomeXchange Consortium ^87^ through the PRIDE partner repository. PX accession number PXD010484.

## ACKNOWLEDGEMENTS

We thank Z. Svindrich and A. Lavanway for help with imaging and image analysis, T. Y. Chang for a generous supply of neurons, W. Wickner for excellent advice on protein purification, M. Pollak for in-put on FSGS, J. McLellan and M. Ragusa for invaluable comments and support, and C. Detteyala for modifying our behavior. This work was supported by NIH R01 GM069818 and R35 GM122545 to HNH, R01 DK088826 to M. Pollak (HNH sub-contract), R35 GM119455 to AK, and P20 GM113132 to the BioMT COBRE.

---

**Figure S1.**
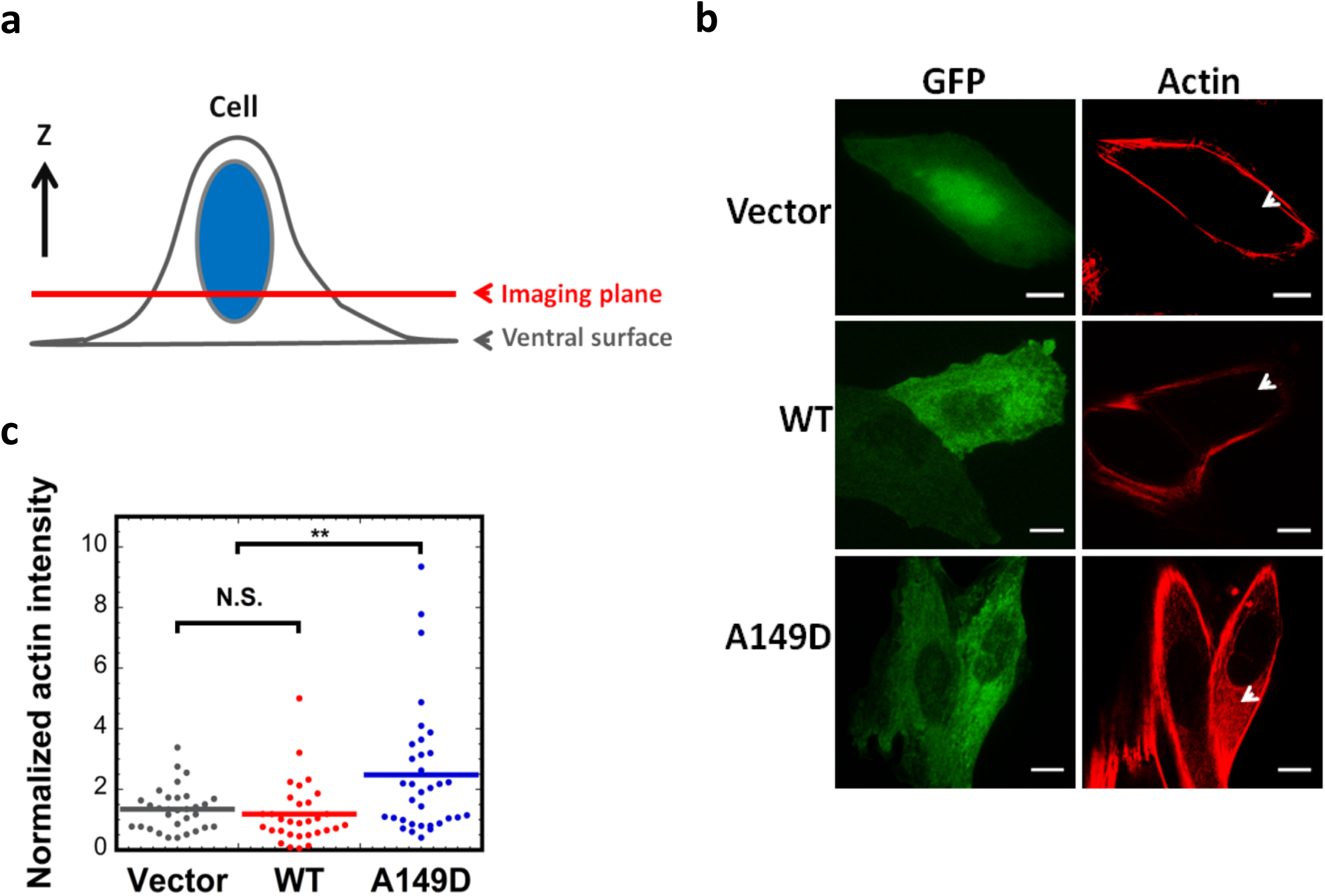
Quantification of cytosolic actin polymerization by exogenously expressed INF2. Confocal microscopy of U2OS cells transfected with GFP alone (vector), GFP-INF2-nonCAAX wildtype (WT) or A149D mutant. Fixed cells stained with TRITC-phalloidin (actin filaments). **(a)** Schematic of confocal imaging plane used. **(b)** Representative GFP and TRITC-phalloidin images. Arrows represent cytosolic regions to be quantified. Scale bar, 10*μ*m. **(c)** Normalized fluorescence intensity of TRITC-phalloidin. Lines represent mean (***Vector: n=31 cells, mean=1.35, STD=0.72; WT: n=31 cells, mean=1.19, STD=1.02; A149D: n=34 cells, mean=2.48, STD=2.14***). *\*\* denotes p* < 0.01 by one-sided student’s t-test. N.S., no significant difference

**Figure S2.**
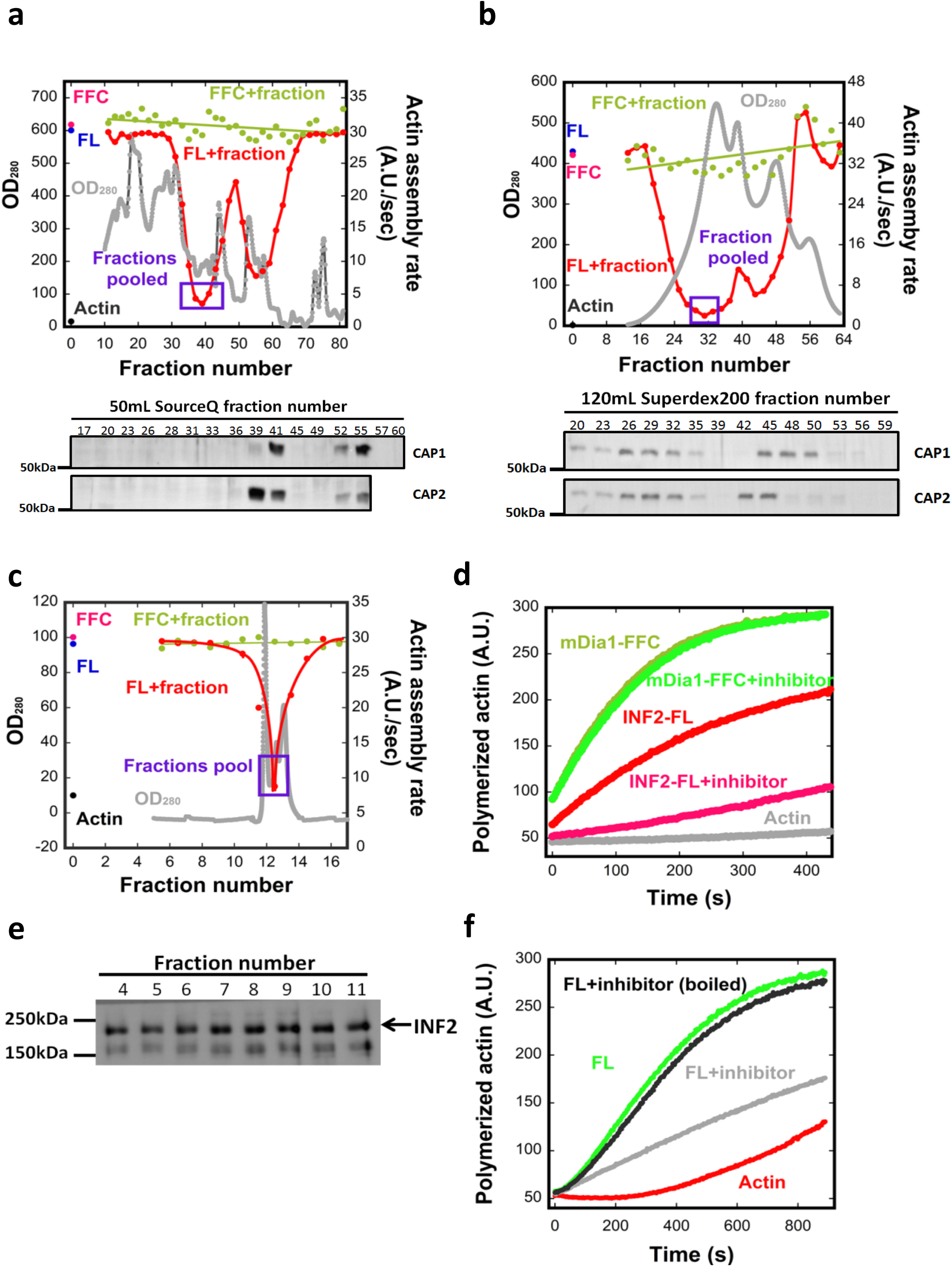
Fractionation of INF2 inhibitory activity by column chromatography. **(a)** Graph showing protein elution profile (OD_280 nm_, gray) from first SourceQ step (pH 7.4) as well as the effect of each fraction on the activities of 20 nM INF2-FL (red) or FFC (green) in pyrene-actin assays. Activity of INF2-FL (blue), INF2 FFC (pink), or actin alone (black) are points on the left of graph. Fractions in purple box are those pooled for subsequent chromatography. Below graph are anti-CAP1 and CAP2 western blots of selected fractions. **(b)** Graph showing protein elution and activity profiles for the Superdex200 size exclusion step. Elution positions of standard proteins (thyroglobulin, ferritin, and γ-globulin) are indicated. Below graph are anti-CAP1 and CAP2 western blots of selected fractions. **(c)** Graph showing protein elution and activity profiles from the second SourceQ (pH 8.8). **(d)** Pyrene actin polymerization assay (2μM actin monomer, 5% pyrene) testing effect of inhibitor (fraction #8 in figure 2e) on 10nM full-length INF2 or 10nM mDia1 FFC. **(e)** Test of GFP-INF2 proteolysis in the presence of inhibitory fractions. Fractions were taken from pyrene-actin assays conducted in Figure 2c, after completion of assay (1200 sec), and probed for INF2 by western blotting. **(f)** Pyrene actin polymerization assays (2μM actin monomer, 5% pyrene) containing 20nM INF2-FL alone (green) or with an inhibitory fraction from final purification step (fraction #8 in figure 2e) that was either boiled (black) or not boiled (gray). Actin alone curve in red.

**Figure S3.**
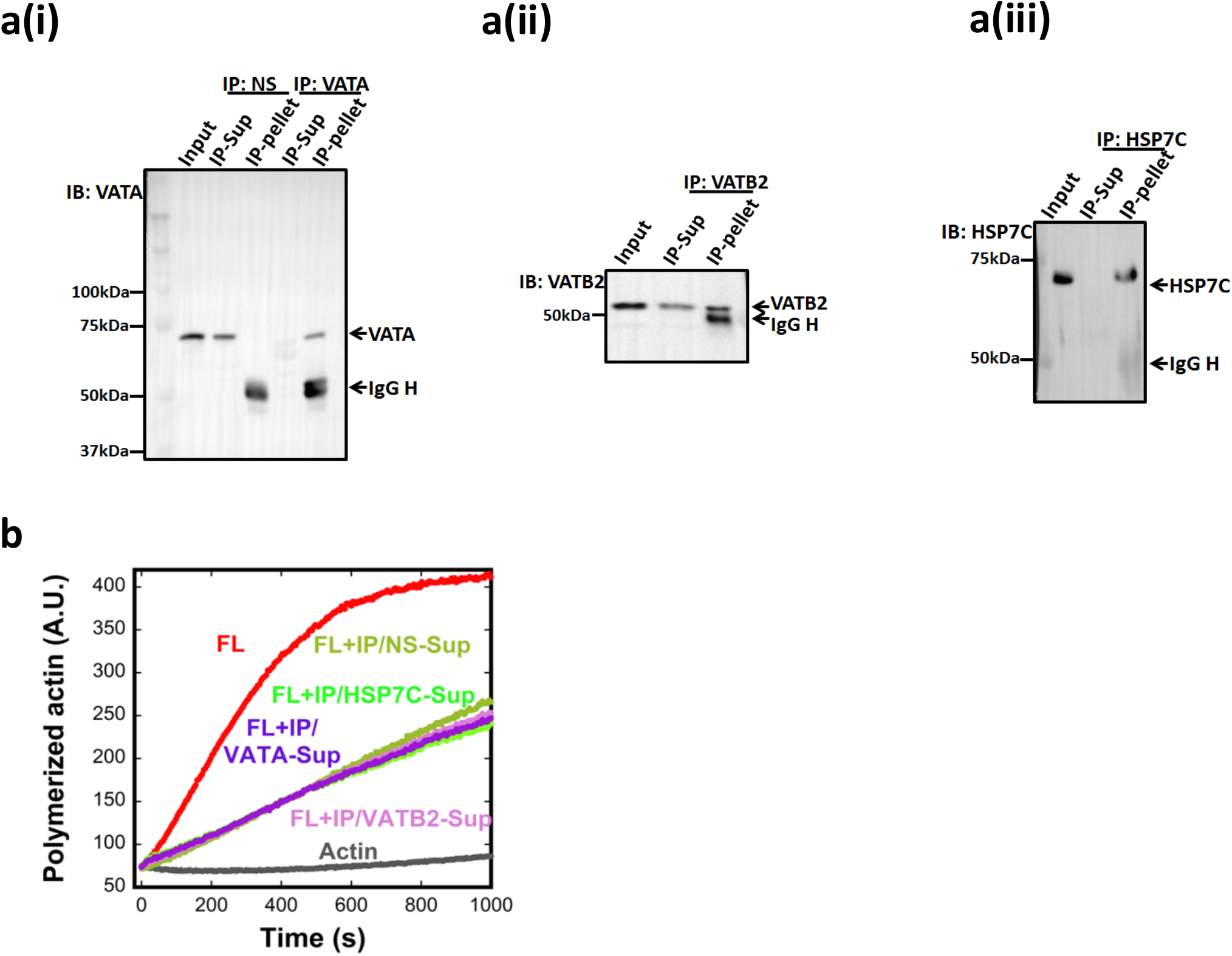
Testing candidate proteins in the BIF for INF2 inhibition by immuno-depletion. **(a)** Immuno-depletion of inhibitor candidates. An INF2 inhibitory fraction from mouse brain (Fraction #38-40 pool from SourceQ#1 (Fig. S2a)) was incubated with either nonspecific antibody control (NS) or the following antibodies: (i) Vacuolar ATPase subunit A (VATA), (ii) Vacuolar ATPase subunit B2 (VATB2, partial depletion), or (iii) Heat shock cognate 71kDa protein (HSP7C). **(b)** Pyrene-actin polymerization assay (2*μ*M actin monomer, 5% pyrene) containing 20nM INF2-FL and supernatant fractions after immuno-depletion. IP/NS-Sup, NS depletion. IP/VATA-Sup, VATA depletion. IP/VATB2-Sup, VATB2 depletion. IP/HSP7C-Sup, HSP7C depletion.

**Figure S4.**
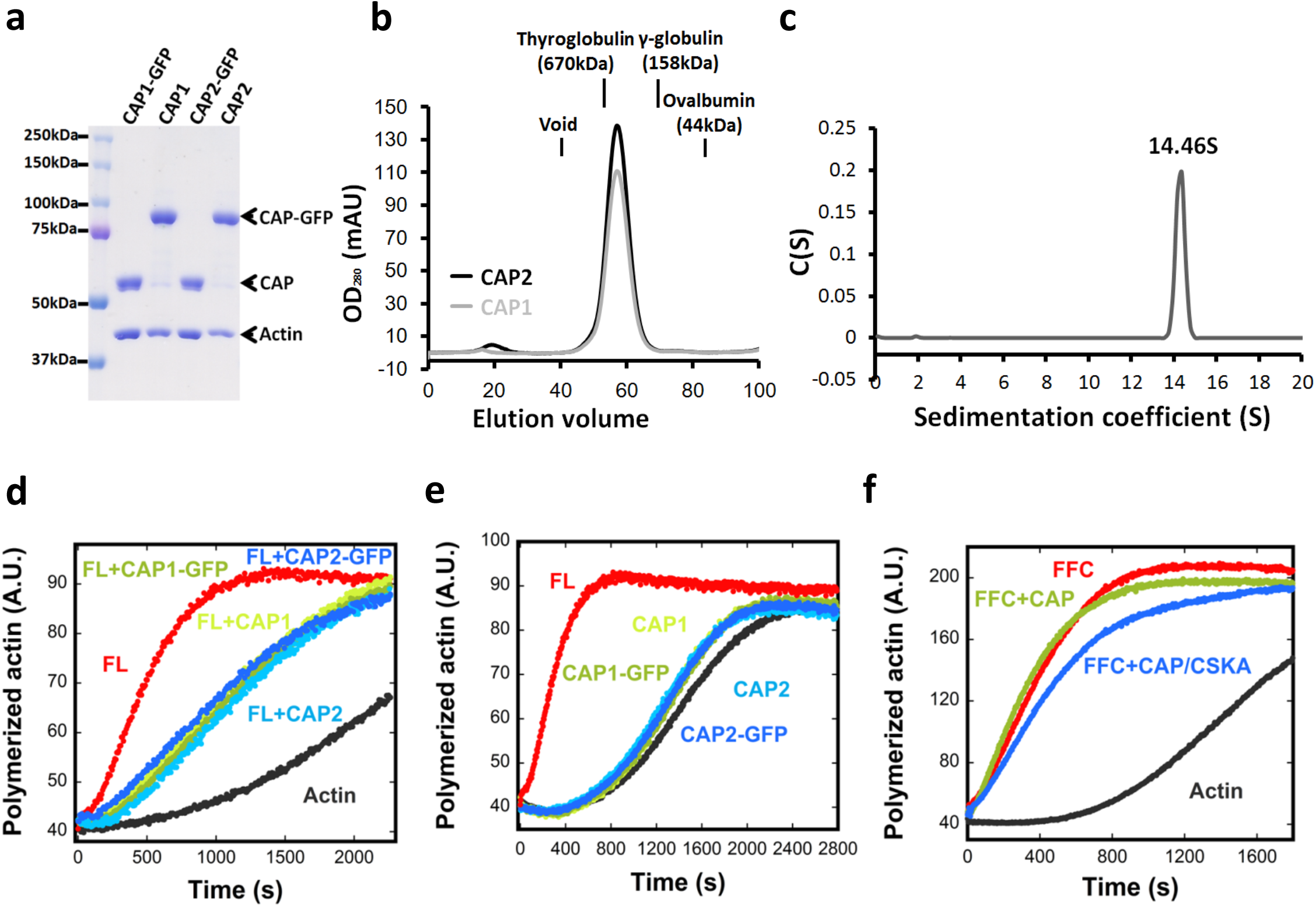
Characterization of recombinant CAP. **(a)** Coomassie stained SDS-PAGE of CAP1 or CAP2 purified from HEK293 cells, either as GFP-fusions or cleaved versions. **(b)** Superdex200 gel filtration chromatography of purified CAP1 or CAP2 (cleaved from GFP). Elution positions of standard proteins (thyroglobulin, ferritin, and *γ*-globulin) indicated. **(c)** Velocity analytical ultracentrifugation of 7*μ*M CAP2-GFP. Mass: 726 kDa (calculated from sedimentation), 83.7 kDa (calculated from sequence). **(d)** Pyrene-actin polymerization assay (2*μ*M actin monomer, 5% pyrene) containing 20nM INF2-FL in the presence or absence of 5*μ*M indicated purified CAP proteins. **(e)** Pyrene-actin polymerization assay (2*μ*M actin monomer, 5% pyrene) in the presence or absence of 5*μ*M indicated purified CAP proteins. **(f)** Pyrene actin polymerization assay (2*μ*M actin monomer, 5% pyrene) containing 20nM GFP-INF2 FFC in the presence or absence of 1*μ*M purified CAP either without actin exchange or exchanged with chicken muscle actin (CAP/CKSA).

**Figure S5.**
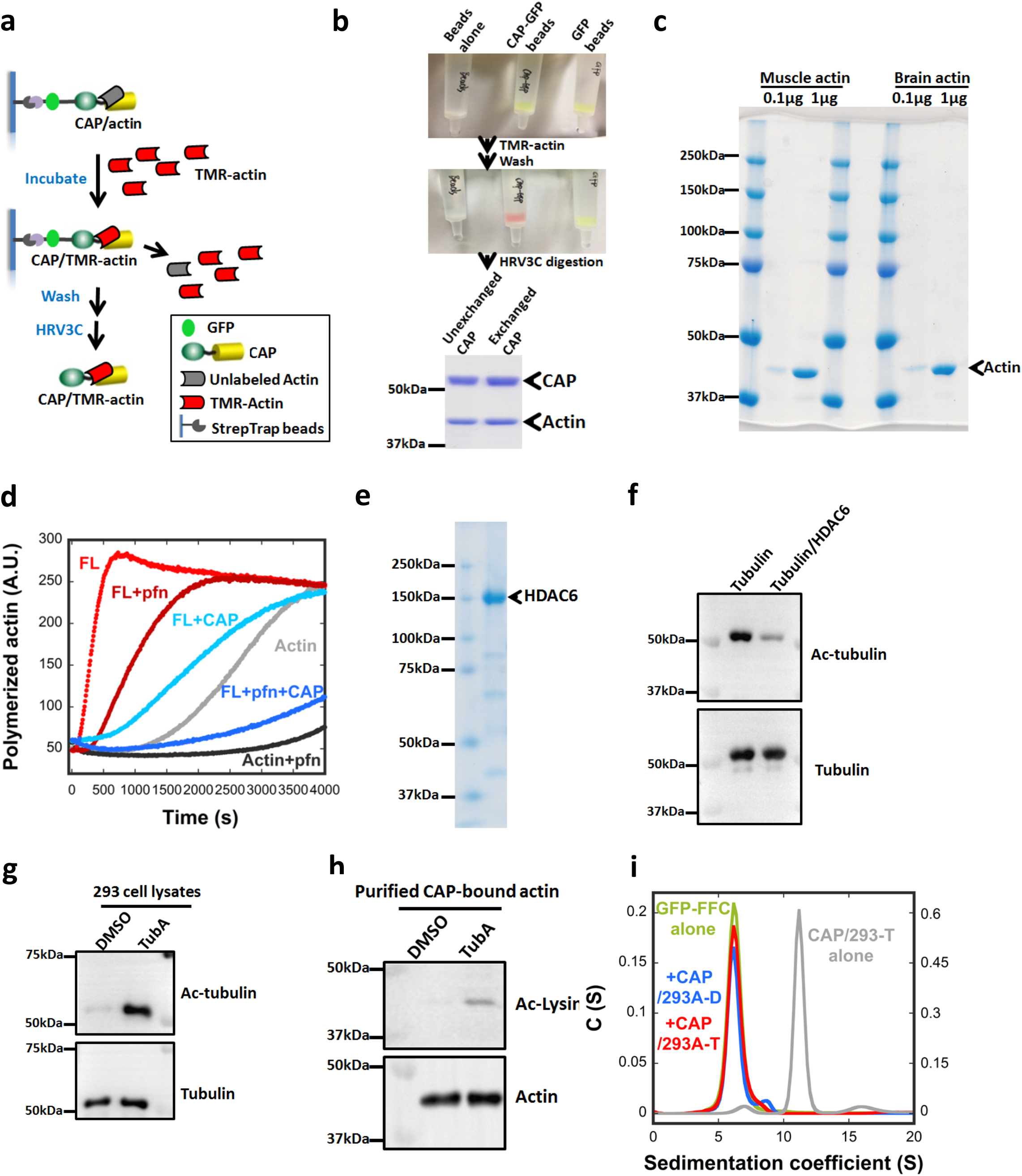
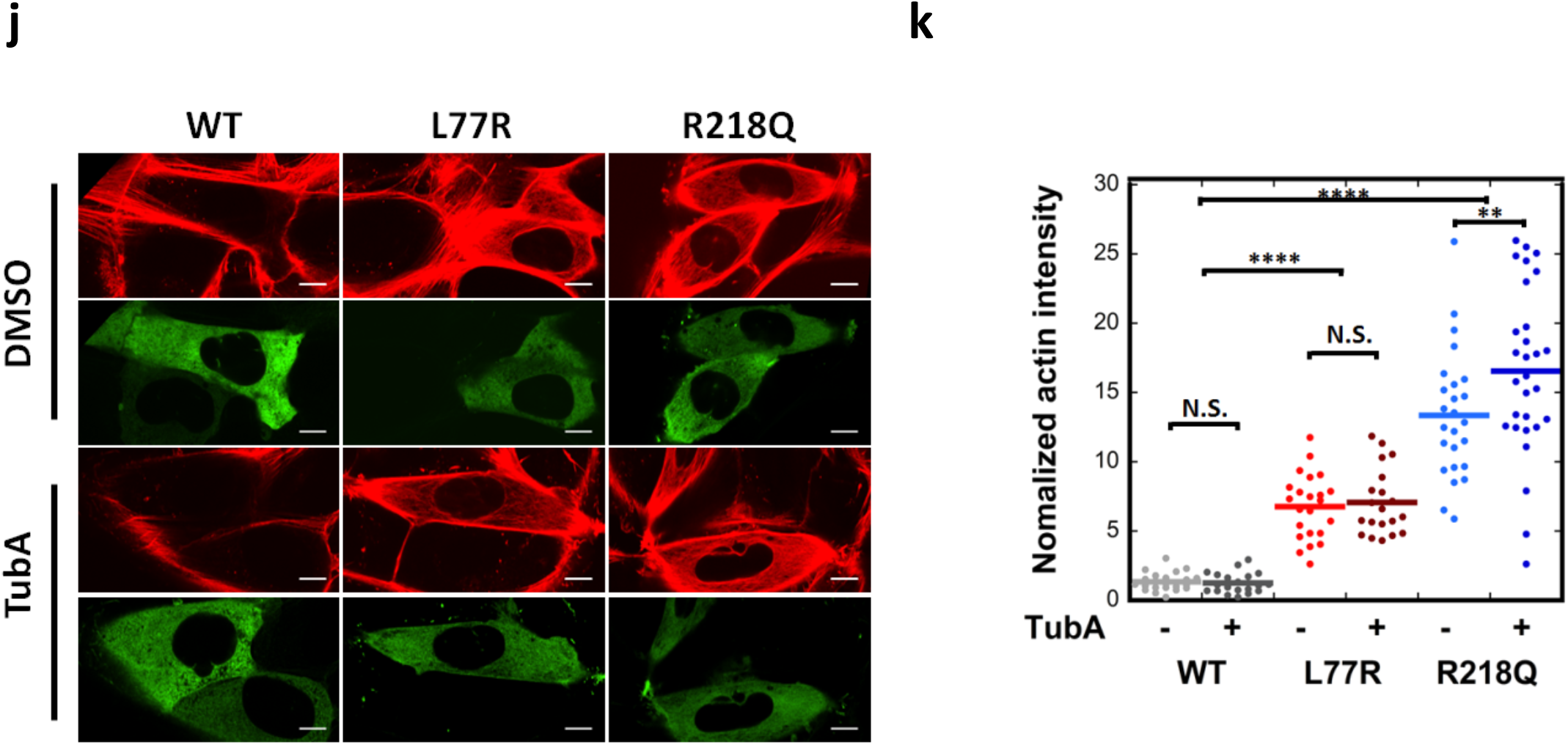
Actin exchange on CAP, and acetylation analysis. **(a)** Schematic of exchange process, in which column-bound CAP/293A is mixed with TMR-actin. CAP/actin is eluted by HRV3C digestion. **(b)** StrepTrap bead chromatographic columns containing no protein (left), CAP1-GFP (middle) and GFP alone (right). Top photo: before mixing beads with TMR-actin. Bottom photo: after mixing with TMR-actin and washing. Coomassie gel shows similar actin content in the HRV3C-eluted complexes after exchange. **(c)** Coomassie gel of purified chicken muscle actin and brain actin. (d) Pyrene-actin polymerization assay (2*μ*M actin monomer, 5% pyrene) containing 20nM GFP-INF2 full-length and 3*μ*M profilin in the presence or absence of 1*μ*M purified CAP2/CSKA. **(e)** Coomassie-stained SDS/PAGE of HDAC6 purified from HEK293 cells. **(f)** Western blot probing tubulin and Ac-tubulin after treatment of purified brain tubulin (10 μM) with purified HDAC6 (200 nM) or buffer for 1hr, 37°C. **(g)** Western blot probing tubulin and Ac-tubulin in 293 lysate after culturing in medium containing DMSO or 50μM Tubastatin A for 1hr prior to lysis. **(h)** Western blot probing actin and Ac-lysine in recombinant CAP2-GFP purified from 293 cells incubated with either DMSO or 50μM Tubastatin A prior to lysis. **(i)** Sedimentation velocity analytical ultracentrifugation of GFP-INF2-FFC (1.8 μM), with or w/o 18μM CAP2/293-D or CAP2/293-T. Mass of GFP-INF2-FFC: 218.3 kDa (from sedimentation), 112.6 kDa (from sequence). **(j)** Representative GFP and TRITC-phalloidin confocal microscopy images showing cytosolic actin polymerization induced by exogenously expressed GFP-INF2 constructs (WT, L77R, or R218Q) in INF2-KO U2OS cells. Cells treated with either DMSO or 50μM Tubastatin A for 1hr followed by fixation and staining with TRITC-phalloidin to label actin filaments. Images are maximum intensity projections of three confocal imaging planes in the middle z region of the cell. Bar, 10 *μ*m. **(k)** Normalized fluorescence intensity of TRITC-phalloidin was quantified from images similar to **(j)**. Lines represent mean. (***WT-DMSO: n=23 cells, mean=1.34, STD=0.64; WT-TubA: n=20 cells, mean=1.25, STD=0.74; L77R-DMSO: n=23 cells, mean=6.76, STD=2.33; L77R-TubA: n=20 cells, mean=7.06, STD=2.38; R218Q-DMSO: n=25 cells, mean=13.35, STD=4.60; R218Q-TubA: n=29 cells, mean=16.55, STD=6.09***). *\*\* denotes p* < 0.01 by one-sided student’s t-test. *\*\*\*\** denotes *p* < 0.0001 by one-sided student’s t-test. N.S., no significant difference

**Figure S6.**
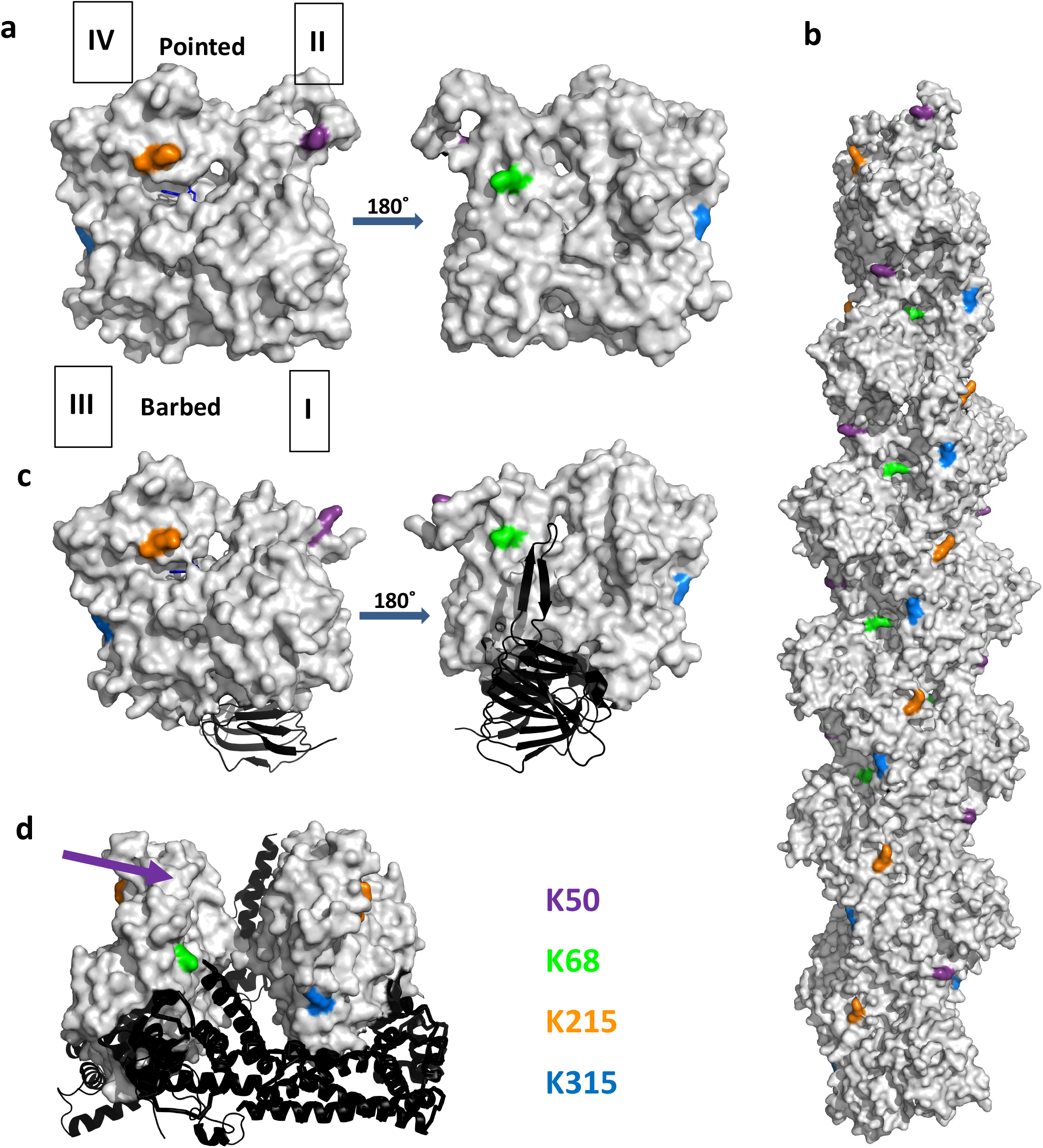
HDAC6-sensitive acetylation sites on actin. Actin in gray with pointed end up in all panels. K50, purple; K68, green; K215, orange; K315, blue. Approximate position of lysine 50 (not resolved in the structure) indicated by purple arrow. Models constructed using Pymol Anaconda2. A) Actin monomer (PDB 4PGK). Actin sub-domains indicated by I, II, III and IV. Structure to right is 180° rotated from the left-hand structure. ATP shown as blue stick figure. B) Actin filament (PDB 3G37). C) Actin bound to CARP domain of mouse CAP1 (PDB 6FM2). Structure to right is 180° rotated from the left-hand structure. ADP shown as blue stick figure. D) Actin bound to FH2 domain of mouse FMNL3 (PDB 4EAH).

